# Pathogen-specific host responses define distinct pneumonia endotypes in the human lung

**DOI:** 10.64898/2026.05.12.724509

**Authors:** Nikolay S. Markov, Marcin Możejko, Vijeeth Guggilla, Małgorzata Łazęcka, Helen K. Donnelly, Alvaro Donayre, Samuel Fenske, Alec Peltekian, Bruno Puczko-Szymański, Paulina Szymczak, Adam Izdebski, Lucy Luo, Karolina J. Senkow, Luisa Cusick, Zhan Yu, Suchitra Swaminathan, Ziyan Lu, Hiam Abdala-Valencia, Duc Phan, Rebecca K. Clepp, Luke V. Rasmussen, Anna Pawlowski, Chiagozie O. Pickens, Nandita R. Nadig, Theresa Walunas, Robert M. Tighe, The Neu-Lung Investigators, Luisa Morales-Nebreda, Catherine A. Gao, Richard G. Wunderink, GR Scott Budinger, Benjamin D. Singer, Alexander V. Misharin, Ewa Szczurek, The NU SCRIPT Study Investigators

## Abstract

Pneumonia is the leading cause of death from infectious disease worldwide. The diagnosis and treatment of patients with pneumonia lag behind other major conditions, relying on syndromic definitions that lack molecular resolution and ignore underlying endotypes. We sought to test the hypothesis that dynamic pathogen-specific host responses in the alveolar space represent distinct pneumonia endotypes linked to different clinical features and outcomes. We prospectively enrolled a cohort of 690 patients (including immunocompromised patients) with known or suspected pneumonia receiving mechanical ventilation in whom the etiology of pneumonia was determined by gold-standard analysis of distal lung fluid obtained by bronchoalveolar lavage (BAL) combined with clinical adjudication. From these patients, we analyzed 792 BAL fluid samples, including 310 serial samples, using flow cytometry (482 patients) and single-cell RNA-sequencing (170 patients; 263 samples, complemented by 9 healthy controls and 25 post-COVID-19 patients, yielding ∼2.4 million single cells across 28 cell types), and extracted daily clinical data from the electronic health record (>15,000 patient-days). We used machine learning models to identify pathogen-specific host responses in the transcriptome of alveolar immune cells that were associated with changes in alveolar cell abundance and clinical features. Our results suggest that therapeutic strategies for pneumonia should be individualized to specific host-pathogen interactions.

## Introduction

While the selection of therapies for many prevalent diseases is increasingly guided by molecular insights into pathogenesis^1^, pneumonia remains an exception. Current clinical guidelines define pneumonia as a syndrome based on respiratory symptoms, physical examination findings, and radiographic evidence of pulmonary infiltrates^2–4^, and recommend antimicrobial and host-directed treatments on this syndromic basis. These guidelines are informed largely by epidemiological studies and animal models, many of which have not been validated in human cohort studies. As a result, guideline-based care applies similar therapy to all hospitalized patients meeting the syndromic definition, necessarily leading to overtreatment. In particular, empirical broad-spectrum antibiotics are recommended regardless of the underlying pathogen, even when a viral etiology is identified, contributing to disruption of the patient microbiome and the emergence of antibiotic-resistant pathogens, a growing public health concern^5^. These limitations may partially explain the persistently high mortality of severe pneumonia, which has remained at 40–50% for more than two decades^6,7^, and may also contribute to the mixed results of clinical trials evaluating host-directed therapies, such as corticosteroids^8,9^, where heterogeneity in the infecting pathogen and the host response are likely to be important.

We sought to determine whether the alveolar immune response during pneumonia is pathogen-specific and is associated with clinical features and outcomes. Accordingly, we conducted a prospective observational study of mechanically ventilated patients with known or suspected pneumonia who underwent clinically indicated gold-standard bronchoalveolar lavage (BAL) sampling to identify pneumonia pathogens^10^. We extracted clinical data from the electronic health record (EHR) and performed multicolor flow cytometry and single-cell RNA-sequencing (scRNA-seq) of BAL fluid. We applied statistical and machine learning methods, ranging from linear factor analysis to transformer-based models, identifying consistent patterns in our data.

We discovered pathogen-specific transcriptomic features of the host response detected by scRNA-seq that are reflected in the cellular composition of the alveolar space as measured by flow cytometry and in clinical features from the EHR. The alveolar host response changed over the course of critical illness, largely driven by changes in the infecting pathogen or response to therapy. Poor outcomes were associated with an exuberant host immune response even when patients were immunocompromised or suffered from repeated infections. These findings suggest pathogen-specific endotypes defined by the host response as potential targets for therapy in patients with severe pneumonia, challenging current treatment guidelines based on syndromic, rather than molecular, definitions. The data portal, including the single-cell object and interactive visualizations, can be explored at https://sqlifts.fsm.northwestern.edu/public/pneumonia-endotypes.

## Results

The syndromic definition of pneumonia is based on the null hypothesis that the clinical presentation of pneumonia is independent of the infecting pathogen^4,11^. To test this definition, we leveraged data and samples from the Successful Clinical Response in Pneumonia Therapy (SCRIPT) study. SCRIPT is a prospective observational cohort study of mechanically ventilated patients, all of whom have undergone at least one BAL procedure for clinical evaluation of known or suspected pneumonia. We included data from the EHR, flow cytometry experiments of BAL fluid to measure cell type abundance, and scRNA-seq experiments in our analysis (**Fig. 1a**).

**Figure 1.**
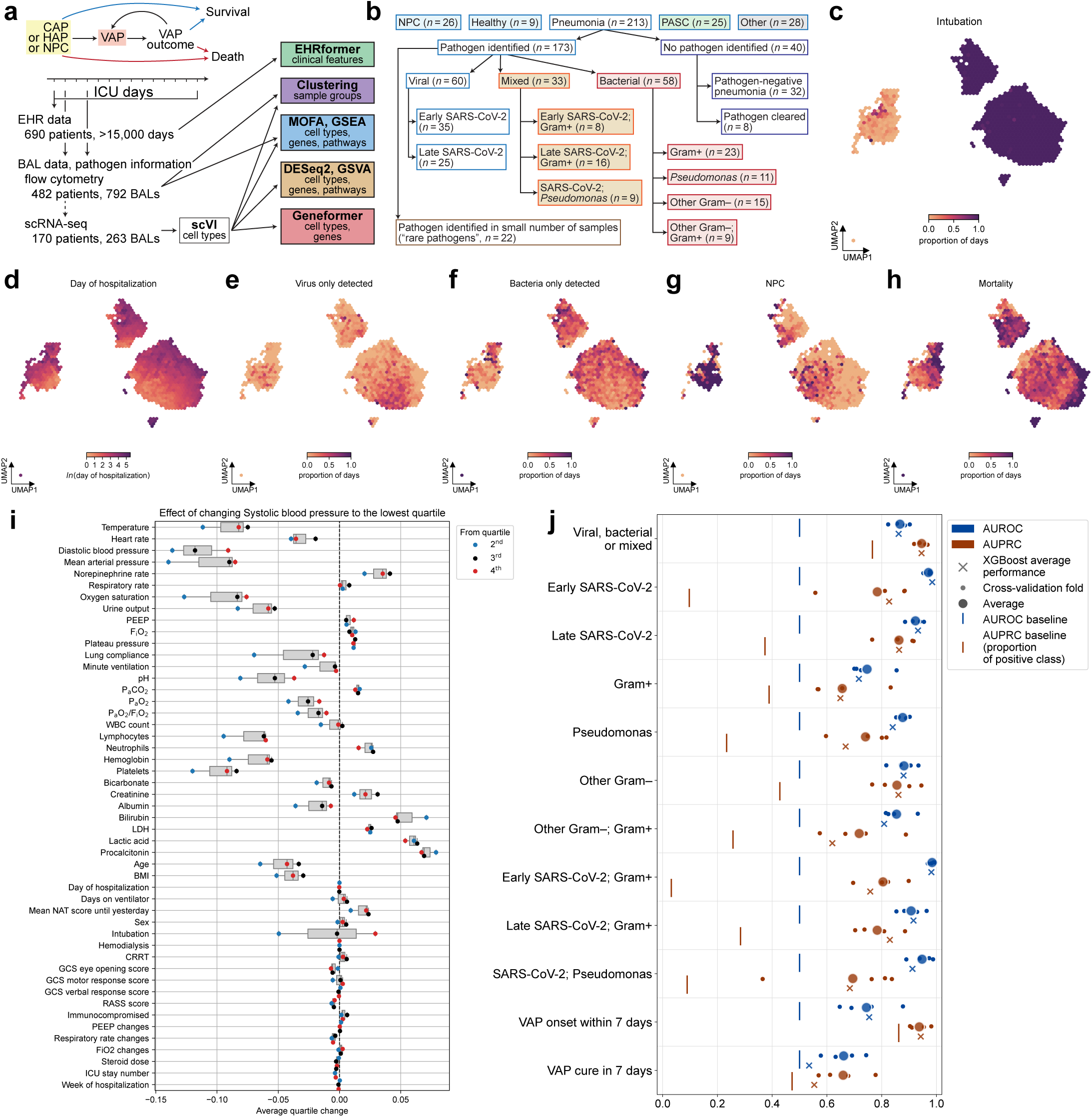
Machine learning applied to data obtained as part of clinical care predicts the pathogen causing pneumonia and VAP onset. **a.** Schematic of the cohort and analytical approach. Community Acquired Pneumonia (CAP), Ventilator Associated Pneumonia (VAP), Intensive Care Unit (ICU), Electronic Health Record (EHR), Bronchoalveolar lavage (BAL), single cell RNA-sequencing (scRNA-seq), Multi-omics Factor Analysis (MOFA), Gene Set Enrichment Analysis (GSEA), Gene Set Variation Analysis (GSVA). **b.** Pathogen groupings in the cohort with the number of BAL scRNA-seq samples. “Pneumonia”: samples clinically adjudicated as pneumonia. “NPC”: samples adjudicated as respiratory failure secondary to a cause other than pneumonia. “Healthy”: healthy volunteers undergoing research bronchoscopy. The adjudicated causes of pneumonia were then subclassified according to the graph. “PASC”: Post-acute Sequelae of SARS-CoV-2 Infection. “Pathogen cleared”: follow-up samples showing pathogen clearance in pneumonia episodes with defined pathogens. **c.** Hexbin plot of UMAP of EHRformer embeddings of ICU patient-day clinical features, colored by the proportion of intubated patient-days in each hexagon. **d.** colored by the day of hospitalization (log scale) in each hexagon. **e.** colored by the proportion of patient-days in which only viral infection was detected in BAL in each hexagon. **f.** colored by the proportion of patient-days in which only bacterial infection was detected in BAL in each hexagon. **g.** colored by the proportion of patient-days from NPC episodes in each hexagon. **h.** colored by the proportion of days from patients with mortality in each hexagon **i.** Effect (average quartile change; x-axis) of simulated perturbation in which the value of systolic blood pressure was set to the lowest quartile, and quartile predictions were obtained from EHRformer for each other clinical feature (y-axis). **j.** Performance of EHRFormer and XGBoost in predicting pathogen classification, VAP onset and cure, as measured by area under receiver operating characteristic curve (AUROC, blue) and area under precision-recall curve (AUPRC, red) on x-axis.

EHR data were available for 690 patients comprising >15,000 patient-days, flow cytometry data were available for 482 patients (792 BAL fluid samples, including 310 serial samples), and scRNA-seq data were available for 170 patients (263 BAL fluid samples) (**Fig. 1a, Extended Data Fig. 1a**). Additionally, we included scRNA-seq data from 9 BAL samples obtained from healthy controls (**Extended Data Table 1**), 25 BAL samples collected from patients with post-acute sequelae of COVID-19 with respiratory symptoms and radiographic abnormalities, both published previously^12^, as well as 4 samples from a cohort of mechanically ventilated patients with severe pneumonia enrolled in an interventional trial (NCT04345614), yielding 301 total scRNA-seq samples comprising ∼2.4 million single cells.

The overall cohort was 59.6% male with a median age of 62 years [IQR 51,71], and 59.3, 20.4, and 19.4% self-reported as White, Hispanic, or Black, respectively. As many patients with severe pneumonia are immunocompromised, we included them in our analysis; they represented 30.1% of the overall cohort. Some patients in this cohort have been described in previous publications (**Extended Data Fig. 1b–d**). Detailed demographic information describing the cohort is found in **Extended Data Table 2**.

The etiology and outcome of all pneumonia episodes were adjudicated by a panel of five critical care physicians using a validated adjudication protocol (**Extended Data Table 2**)^13^. From 927 BAL samples collected from 687 patients, the panel identified a respiratory pathogen as the cause of a pneumonia episode in 619 samples (67%) from 452 patients. In 194 (21%) of the BAL samples, the panel adjudicated the episode as pneumonia in the absence of an identified pathogen (“pathogen-negative pneumonia”). In 114 (12%) of the BAL samples, the panel adjudicated the episode as respiratory failure secondary to a cause other than pneumonia, a group we refer to as “non-pneumonia controls” (NPC).

As our study overlapped with the COVID-19 pandemic, our samples were enriched in patients with SARS-CoV-2 pneumonia. Within the group of patients with complete clinical, flow cytometry, and scRNA-seq data available, we defined major pathogen subgroups as the cause of the pneumonia, defined by our clinical adjudication procedure^13^, for subsequent comparisons (**Fig. 1b, Supplementary Table 1**). These subgroups included infection types that were common enough to allow statistical comparisons (group size of at least 8 samples).

To identify pathogen-specific features of the host response to pneumonia, we analyzed our data using newly developed and existing data science tools. These included a novel transformer-based model for EHR records (EHRformer)^14^, Multi-Omics Factor Analysis (MOFA^15^), clustering of samples, differential gene expression analysis (DESeq2^16^), a transformer-based foundation model for scRNA-seq data that was fine-tuned on our data (Geneformer^17^), and pathway enrichment analysis (GSEA^18^ and GSVA^19^) (**Fig. 1a**).

### EHR data predict the infecting pathogen in patients with suspected pneumonia

An exploratory analysis of the overall structure of the data suggested pneumonia endotypes across modalities that were evident from sample-level correlations between pathogens, flow cytometry, and scRNA-seq data with clinical features, including mortality (**Extended Data Fig. 2**). We began by exploring the EHR data using transformer- and tree-based models^14,20^. We asked the models to predict the pathogen causing the pneumonia episode defined by BAL fluid results and clinical adjudication^13^, which we extrapolated to patient-days following BAL procedures. In addition, because a prevalent hypothesis predicts that viral pneumonia predisposes the host to subsequent bacterial infection^21^, we asked the models to predict the development or cure of ventilator-associated pneumonia (VAP). We extracted 50 clinical features from the EHR for every day in the ICU for each patient, and made these data available on Physionet for research^22^ (**Extended Data Table 2**). These features are routinely measured in the ICU and include dichotomous variables (e.g., receipt of hemodialysis or continuous renal replacement therapy [CRRT], extracorporeal membrane oxygenation [ECMO], and intubation for mechanical ventilation) and continuous variables (e.g., vital signs, mechanical ventilator parameters, laboratory test results). We also included selected medication dosages as continuous variables, including vasopressor infusion rates used to maintain arterial blood pressure (summarized as norepinephrine dose), steroid doses (summarized as the equivalent anti-inflammatory dose of hydrocortisone), and the relative pathogen spectrum of the antibiotics administered on each day of their ICU stay (summarized by the Narrow Antibiotic Treatment Score, which ranges from more narrow than guideline recommended therapy to more broad than guideline-recommended therapy for community-acquired pneumonia^23^).

To model these clinical features, we developed EHRformer, a four-layer transformer-based model for tabular input, and pre-trained multiple models on subsets of our EHR data depending on the predictive task^14^. We explored the pre-trained EHRformer embeddings of patient-days to understand trends in the EHR data captured by the model. The two major clusters of patient-days in a UMAP visualization of the embeddings corresponded to intubation status (**Fig. 1c**). Other axes of variation in the embeddings included the duration of hospitalization (**Fig. 1d**) and the severity of illness as measured by the Sequential Organ Failure Assessment (SOFA) score (**Extended Data Fig. 3a**). Patient-days for which only viral pathogens were detected by PCR fell in a different region of the embedding space than patient-days with only bacterial pathogens detected by PCR or culture, with detection extrapolated as pathogen results (**Fig. 1e, f**). NPC patient-days partially overlapped with the non-intubated cluster and partially with the bacteria-infected rather than virus-infected embedding space (**Fig. 1g**). Patient-day embeddings from patients with early SARS-CoV-2 separated from patients with late SARS-CoV-2 (**Extended Data Fig. 3b, c**). Hospital mortality/died (defined as death, discharge to hospice or lung transplantation for refractory respiratory failure during the index hospitalization) was more intermixed and less visually separated, reflecting the severe illness within the cohort (**Fig. 1h)**.

To assess whether EHRformer captured clinically meaningful relationships among features, we performed simulated perturbations in the latent space of EHRformer, setting individual feature values to the highest or lowest quartile in the dataset and examining the changes in other features reconstructed by the model (**Fig. 1i, Extended Data Fig. 3d**). These perturbations were clinically interpretable and biologically meaningful, validating that the model correctly learned the underlying pathophysiology. For example, when systolic blood pressure was set to the lowest quartile, EHRformer predicted higher doses of norepinephrine, higher levels of lactic acid, and higher levels of secondary organ injury (serum creatinine and bilirubin) (**Fig. 1i**). When positive end-expiratory pressure (PEEP) was set to the highest quartile, EHRformer predicted higher F_i_O_2_ and respiratory rate and lower alertness (**Extended Data Fig. 3d**).

We fine-tuned EHRformer to distinguish patient-days labeled as NPC from any viral, bacterial or mixed pneumonia, and from each of 9 pathogen groups individually, and additionally fine-tuned it to predict VAP onset within 7 days and VAP cure within 7 days. We compared performance against XGBoost, a tree-based algorithm commonly applied to tabular clinical data, and against a random baseline (**Extended Data Fig. 3e–h**, **Supplementary Table 2**). EHRformer outperformed XGBoost for the majority of tasks as measured by area under the receiver operating characteristic curve (AUROC; random = 0.5) and area under the precision-recall curve (AUPRC; random = fraction of positive class), the latter being more appropriate for class-imbalanced tasks (**Fig. 1j**, **Supplementary Table 3**). Across pathogen-specific tasks, EHRformer AUPRC was several-fold higher than the random baseline, with the highest performance for distinguishing NPC from any pathogen (AUPRC 0.95) and for late SARS-CoV-2 and other gram-negative bacteria (AUPRC 0.74–0.95). Both models performed well for VAP onset prediction (AUPRC 0.94), though performance for VAP cure was more modest (AUPRC 0.66 and 0.55 for EHRformer and XGBoost, respectively), suggesting that resolution of VAP is less legible from routine clinical data alone.

To identify the clinical features most informative for pathogen identification and VAP prediction, we computed SHapley Additive exPlanations (SHAP) values for XGBoost and extracted attention weights for all 50 clinical features from the transformer layers of the EHRformer models. We summarized them for patient-days either correctly classified as NPC, each of the pneumonia pathogen groups, or each VAP prediction task (**Extended Data Fig. 3f–h**). SHAP values for XGBoost were dominated by age, BMI, and days on ventilator, while attention scores for EHRFormer were driven by physiologic measurements and laboratory values associated with pathogen detection or illness severity, including lactic acid levels, level of PEEP, total white blood cell count (WBC), norepinephrine dose, and urine output. The attention patterns varied between correctly classified NPC and pathogen-specific samples, suggesting that the model differentiates features based on the subgroup class (**Extended Data Fig. 3g**). For example, the attention to procalcitonin and mean arterial pressure was more pronounced in pathogen-specific tasks compared with NPC, highlighting the relevance of these features in distinguishing between different etiologies of pneumonia and respiratory failure^24^. The divergence between XGBoost and EHRformer feature importance suggests that transformer-based modeling captures different and more physiologically grounded aspects of the clinical signal.

In summary, our analysis of EHR data revealed pathogen-specific features of the alveolar host response to severe pneumonia that might be used alongside molecular tests to guide antimicrobial and host-directed therapies.

### The cellular composition of the alveolar space reflects pathogen-specific host responses

Before the COVID-19 pandemic, we reported that a relative paucity of neutrophils (<50% of BAL fluid immune cells) in the alveolar space has a negative predictive value of >90% for pneumonia from any pathogen^25^. In patients with SARS-CoV-2 pneumonia, however, we observed relatively low levels of neutrophils in BAL samples collected within 48 hours of intubation^26^. These results suggest the relative abundance of inflammatory cells in the alveolar space reflects pathogen-specific features of the host response to pneumonia. To test this hypothesis, we developed a customized flow cytometry panel to detect the major inflammatory cell populations in the alveolar space^12,26^. We used a marker of mature alveolar macrophages, CD206 (encoded by *MRC1*), to distinguish monocyte-derived alveolar macrophages that had recently differentiated from recruited monocytes (CD206^lo^) from more mature (CD206^hi^) alveolar macrophages^12^.

In BAL fluid from healthy controls, most of the sampled cells were alveolar macrophages (78%) followed by lymphocytes (17%), while neutrophils and other granulocytes combined accounted for only 5% of cells^12^. Samples from NPC and patients with pneumonia were clustered based on the relative abundance of cell populations detected by flow cytometry (**Extended Data Table 3**, **Supplementary Table 4**). We named these clusters according to the most abundant cell types (**Fig. 2a, b**). We identified associations between the clusters and clinical features, testing additionally for enrichment in samples belonging to different adjudicated pathogen groups (**Fig. 2c** and **Extended Data Fig. 4a, b**). Many of these associations have been identified previously, suggesting this clustering procedure captures meaningful associations^26,27^. For example, the *Neutrophil+* and *Neutrophil+Monocyte* clusters were associated with an adjudicated bacterial pathogen^25^, while the *T cell+Treg* and *Monocyte+* clusters were associated with early SARS-CoV-2 pneumonia (**Fig. 2c**). The *Mature macrophage+* cluster, a characteristic of normal lung, was associated with samples from NPC (**Fig. 2c**).

**Figure 2.**
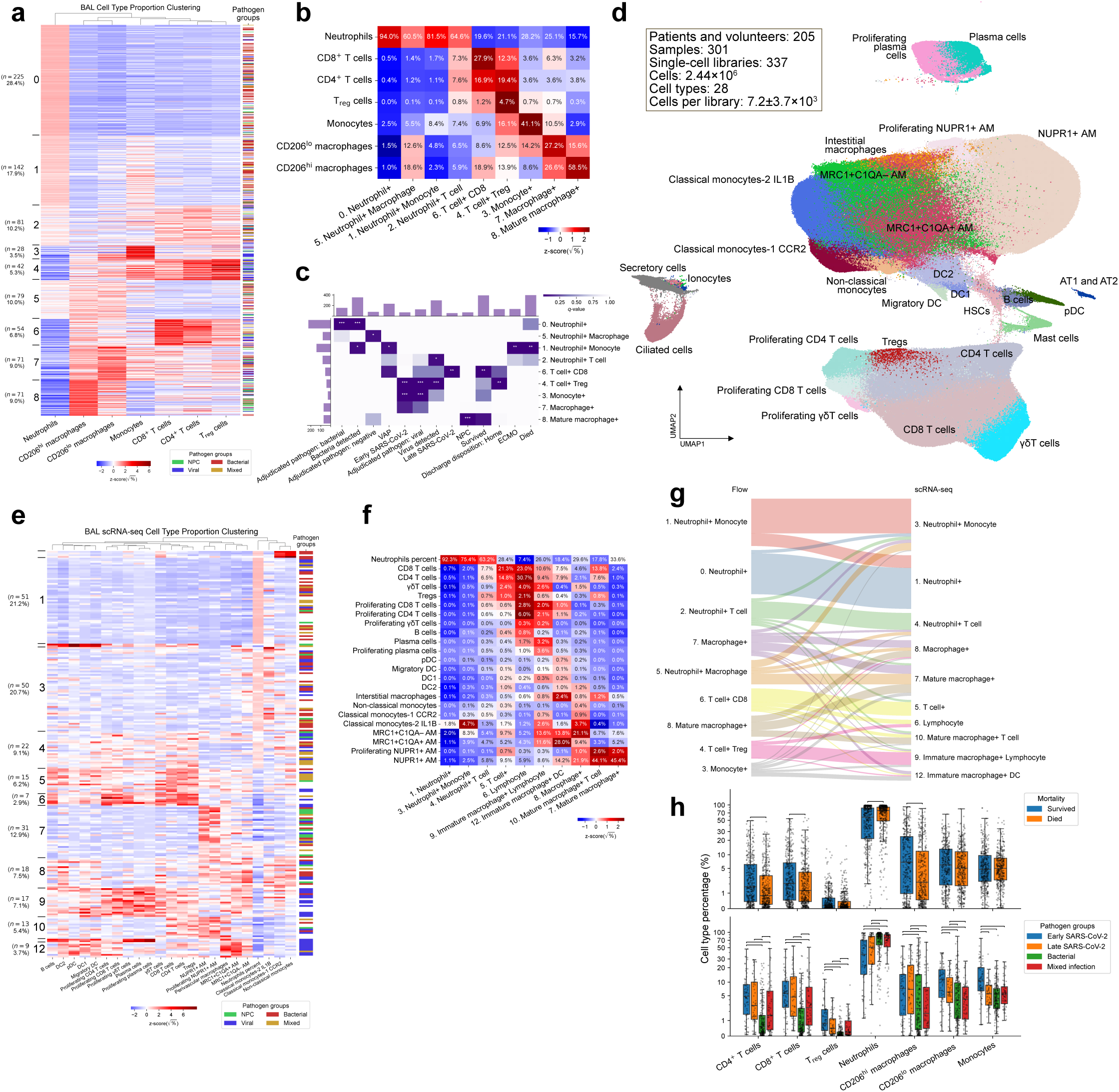
Pathogen specific differences in the cellular composition of the alveolar space. **a.** Heatmap of relative abundance of different cell populations detected by flow cytometry (columns) in samples (rows) ordered by sample clusters (left margin, including cluster number, number of samples and percentage of samples in cluster). CD206^high^ and CD206^low^ macrophage: alveolar macrophages with relatively high or low surface staining for CD206, Treg: regulatory T cells. Pathogen groups include non-pneumonia controls (NPC), or an episode of pneumonia secondary to a virus, bacteria, or both virus and bacteria (viral, bacterial, mixed, respectively). **b.** Normalized average percentage of the cell populations (rows) in each cluster (columns) as detected using flow cytometry. Clusters were named based on one or two most abundant cell populations. **c.** Associations between sample clusters determined by flow cytometry (rows) and pathogen groups or selected clinical features (columns). Pathogen groups and clinical features with significant associations are shown (right-tailed hypergeometric tests with FDR correction; * *q*-value < 0.05, ** *q*-value < 0.01, *** *q*-value < 0.001). “Adjudicated pathogen”: expert adjudication of pathogen cause of pneumonia episode (viral, bacterial, mixed, negative [pneumonia in the absence of identified pathogen]). “Bacteria detected” or “Virus detected”: a positive PCR or culture of BAL for a bacterium or virus, respectively. “VAP”: ventilator-associated pneumonia. “Early SARS-CoV-2” or “Late SARS-CoV-2”: BAL sample positive for SARS-CoV-2 collected ≤ or > 48 hours after intubation, respectively. “NPC”: non-pneumonia control. “VAP cure in 7 days”: clinical adjudication of VAP episodes resolved within 7 days. “ECMO”: the requirement for extracorporeal membrane oxygenation. A full list of clinical features with significant associations is provided in **Extended Data** Fig. 4a. **d.** UMAP projection of scRNA-seq data for this cohort. This includes 263 samples from 170 patients with known or suspected pneumonia, 9 healthy control subjects, 25 samples from 24 patients with persistent respiratory symptoms more than 60 days after SARS-CoV-2 pneumonia (PASC), and 4 samples from 2 patients enrolled in a clinical trial of Auxora. Monocytes and alveolar macrophages are annotated along a continuum of maturation: Classical monocytes-1 CCR2 (circulating classical monocytes) and Classical Monocytes-2 IL1B (alveolar monocytes), MRC1+C1QA– AM (immature alveolar macrophages), MRC1+C1QA+ AM (maturing alveolar macrophages), NUPR1+ AM (mature alveolar macrophages). DC - dendritic cell, pDC - plasmacytoid dendritic cell. AT1 and AT2 - alveolar type 1 and 2 epithelial cells, respectively. **e.** Relative abundance of different cell populations detected by scRNA-seq (columns) in samples (rows) ordered by sample clusters (left margin, including cluster number, number of samples and percentage of samples in cluster). Clusters 0, 2 and 11, which had fewer than 5 samples, are not labeled. **f.** Normalized average percentage of the cell populations (rows) in each cluster as detected using scRNA-seq (columns). Clusters were named based on one or two most abundant cell populations. Clusters 0, 2 and 11, which had fewer than 5 samples, are not shown. **g.** Sankey plot comparing assignment of samples to cellular abundance clusters using flow cytometry (left) and scRNA-seq (right) for a common subset of samples (*n* = 233). **h.** Percentages of cell type populations as detected by flow cytometry, split by mortality (top) and pathogen groups (bottom). Brackets denote significant comparisons (*q*-value < 0.05, two-sided Mann-Whitney U tests with FDR correction).

BAL fluid enrichment in T cells (*T cell+Treg* and *T cell+CD8* clusters) was associated with favorable outcomes. For example, the *T cell+Treg* cluster was associated with discharge disposition to home (**Fig. 2c**). We and others have reported that regulatory T (Treg) cells promote recovery after lung injury in mouse models, and clinical trials of Treg cell therapies for patients with ARDS have shown promise^28^. In contrast, a relative abundance of BAL fluid monocytes and neutrophils (*Neutrophil+Monocyte*) was associated with poor outcomes including mortality, receipt of ECMO support, and VAP (**Fig. 2c**). Combined with the ability of EHRformer to predict VAP, this finding lends support to a hypothesis from animal models and clinical reports that viral pneumonia enhances the risk of secondary bacterial infection^21,29,30^ .

To perform a more granular investigation of cell type composition, we turned to scRNA-seq data and resolved 28 cell types, including monocytes, macrophages, dendritic cells, T cell subsets, B cells and plasma cells, and a small number of airway and alveolar epithelial cells (**Fig. 2d, Extended Data Fig. 4c, Extended Data Table 4**). Monocytes and macrophages formed a continuum of transcriptional states, reflecting the differentiation of classical monocytes recruited from the circulation into progressively more mature alveolar macrophages during injury, as described in murine models^31–33^. We detected classical monocytes typically seen in the circulation (“Classical monocytes-1 CCR2”), which likely represents bleeding into the alveolar space and detection of peripheral blood monocytes in BAL fluid. Upon migration into the alveolar space, classical monocytes lose their expression of *CCR2* and upregulate *IL1B* (“Classical monocytes-2 IL1B”). *MRC1* is among the hundreds of genes that increase as monocytes mature into alveolar macrophages, followed later by *C1QA*. This sequential expression is represented by our nomenclature: “MRC1+C1QA– AM” of immature and “MRC1+C1QA+ AM” of maturing alveolar macrophages, respectively. Mature alveolar macrophages (“NUPR1+ AM”) are characterized by expression of genes associated with homeostatic processes, lipid metabolism, and adaptation to long-term persistence in the alveolar space. They are the dominant population in healthy volunteers^12,34^.

We estimated the abundance of immune cell populations in the alveolar space within each sample and identified associations between alveolar cell type abundance, pathogens, and clinical features using the same procedures as for the flow cytometry data (**Fig. 2e–h, Extended Data Fig. 4d–f**). Because the collection and storage protocol used to archive SCRIPT specimens excluded neutrophils from our scRNA-seq data, we included their abundance from flow cytometry data, as we have described previously^12^. We could not test for associations with VAP onset in these data because we did not have sufficient scRNA-seq samples collected from patients before VAP developed. We sought to confirm associations we identified using flow cytometry in the subset of samples for which scRNA-seq data were available. We were able to reproduce 7/21 associations (FDR q < 0.05) (**Extended Data Fig. 4e**). These included associations between “Neutrophil+ Monocyte” with mortality and “Mature macrophage+” with NPC (**Extended Data Fig. 4e**). The higher resolution of the single cell analysis identified associations between “Immature macrophage+ Lymphocyte” and “Immature macrophage+ DC” with early SARS-CoV-2 pneumonia. These associations might result from SARS-CoV-2-induced death of alveolar macrophages, as we have observed and others have reported in animal models of viral pneumonia^26,29^. Consistent with this hypothesis, SARS-CoV-2 transcripts were identified in epithelial cells and immature alveolar macrophages. (“MRC1+C1QA- AM”) (**Extended Data Fig. 4g**).

### The transcriptome of alveolar immune cells is pathogen-specific

To discover major factors of variation in both abundances of major immune cell types derived from scRNA-seq and pseudobulk transcriptomic profiles of those cell types, we provided them as inputs to Multi-Omics Factor Analysis (MOFA) (**Fig. 3a**). MOFA factors, indexed by samples, represent reduced data dimensions and tend to capture major linear trends behind the data. The weight vectors (one per factor), indexed by features, link the factors to the underlying molecular processes within each cell type.

**Figure 3:**
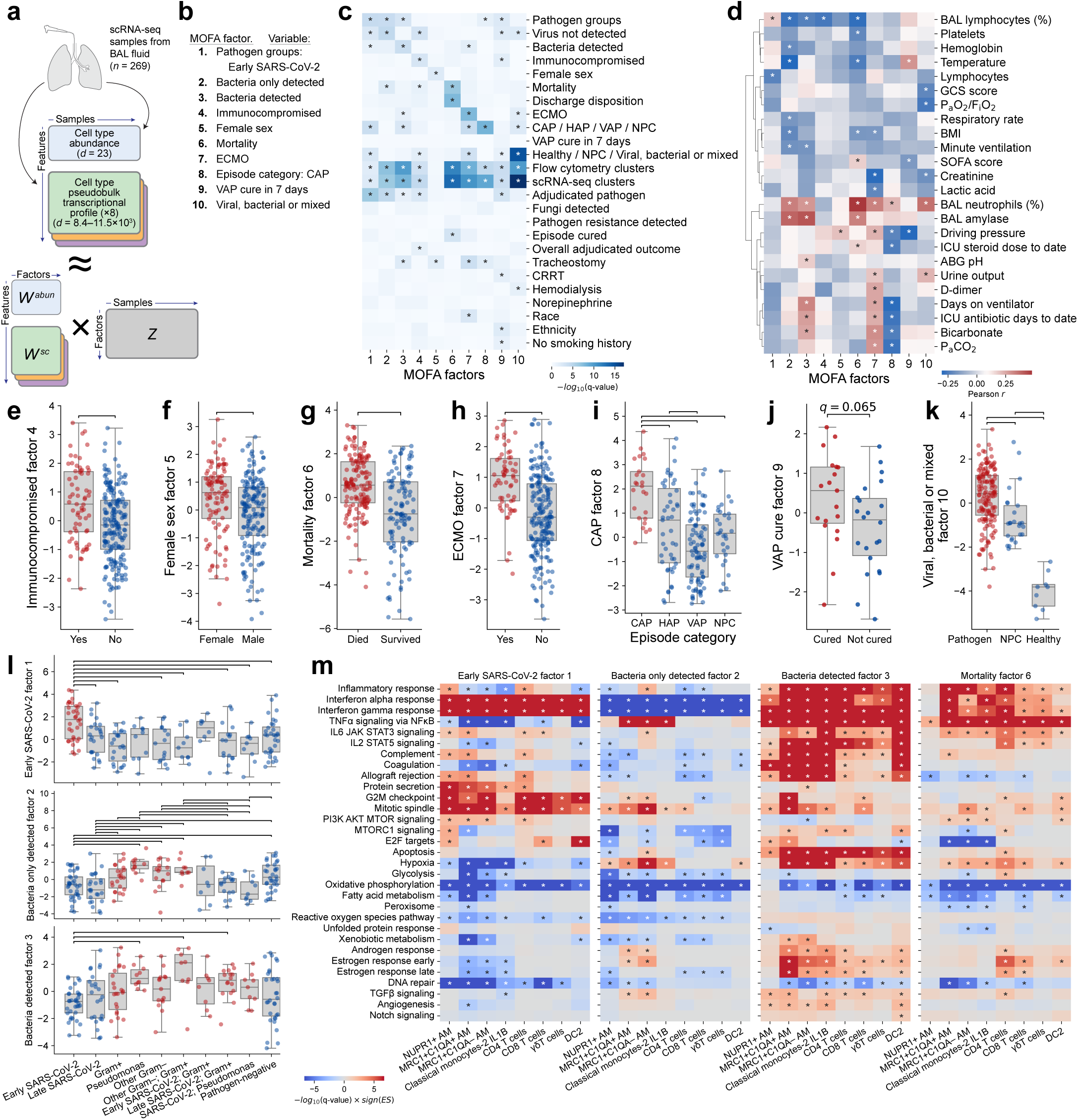
Major axes of variation behind cell type-specific RNA-seq and cell type abundance data. **a.** MOFA analysis overview. We organized the data into 9 views, with each view given by a samples × features matrix, for 269 samples and for the number of features specific for each view, with a total of 80,788 features across all views. The cell type abundance view had one feature per each cell type, while the views corresponding to cell-type gene expression had the number of features equal to the number of genes robustly expressed for each cell type. **b.** Rotations of factors performed to encourage associations with 10 predefined features. **c.** Associations between 10 factors (columns) and pathogen categories and categorical clinical features (rows). * indicates significant association as determined by two-sided Mann-Whitney U test (for categories with 2 levels) or ANOVA test with FDR correction. **d.** Association between 10 MOFA factors (columns) with continuous clinical features (rows). * indicates significant association as determined by Spearman correlation with FDR correction. **e.** Values of “immunocompromised factor 4” for samples from patients classified as immunocompromised (yes) compared to those who were not (no). Bar indicates two-sided Mann-Whitney p-value < 0.05. **f.** Values of “female sex factor 5” for samples from patients with female compared to male sex. Bar indicates two-sided Mann-Whitney p-value < 0.05. **g.** Values of “mortality factor 6” for samples from patients who died compared to those who survived their hospital stay. Bar indicates two-sided Mann-Whitney p-value < 0.05. **h.** Values of “ECMO factor 7” for samples from patients who received ECMO at the time of sampling compared to those who did not. Bar indicates Mann-Whitney p-value < 0.05. **i.** Values of “community acquired pneumonia (CAP) factor 7” for samples from patients with a pneumonia episode adjudicated as CAP, Hospital Acquired Pneumonia (HAP), Ventilator Associated Pneumonia (VAP), or Non Pneumonia Control (NPC). Bar indicates FDR-corrected two-sided Mann-Whitney q-value < 0.05. **j.** Values of “VAP cure factor 9” for samples from patients whose VAP episode was adjudicated as cured or not cured. Differences were not significant. **k.** Values of “viral, bacterial or mixed factor 10” for samples with viral, bacteria or mixed pathogen (Pathogen), NPC, or a healthy control. Bar indicates FDR-corrected two-sided Mann-Whitney q-value < 0.05. **l.** Values of “early SARS CoV-2 factor 1” (top), “bacteria only detected factor 2” (middle), and “bacteria detected factor 3” (bottom) for pathogen groups (x-axis). Bar indicates FDR-corrected two-sided Mann-Whitney q-value < 0.05. **m.** GSEA enrichment of hallmark gene sets (rows) in gene lists ranked by MOFA weights per each cell-type view (columns; Extended Data Fig. 5a) for factors “early SARS-CoV-2 factor 1”, “bacteria only detected factor 2”, “bacteria detected factor 3” and “mortality factor 6”. * indicates significant enrichment using GSEA after FDR correction.

We fitted a MOFA model with 10 factors, achieving an average of 30% explained variance for the dataset (**Extended Data Fig. 5a**). Gene expression and to a lesser extent cell type abundance in multiple cell populations contributed to the variance identified by MOFA, with most variance explained for macrophage subtypes (**Extended Data Fig. 5a**). We applied a factor rotation^35^ to encourage associations with samples from pathogen group categories: early SARS-CoV-2 for factor 1; bacteria only detected for factor 2; and samples with bacteria detected for factor 3 (**Fig. 3b, Extended Data Fig. 5b**). We also encouraged associations with the following: demographic variables known to confer an increased or reduced risk for pneumonia (immunocompromised state for factor 4 and female sex for factor 5); outcome variables (mortality for factor 6 and receiving ECMO for factor 7); guideline-based definitions of CAP for factor 8; VAP cure in 7 days for factor 9; and BAL from viral, bacterial or mixed pathogen group for factor 10 (**Fig. 3b, Extended Data Fig. 5b**). The resulting rotated MOFA factors are hereafter referred to simply as factors.

We queried the resulting factors for associations with other variables within our clinical data (**Fig. 3c,d**) and confirmed the rotation procedure by inspecting factor distributions across clinical categorical variables (**Fig. 3e–l**). Despite rotational encouragement toward VAP cure by day 7, no factor with a meaningful association emerged, suggesting either insufficient signal in the data or a non-linear relationship not captured by MOFA (**Fig. 3c**). We attempted additional rotations toward the more specific pathogen groups (e.g., *Pseudomonas*), but failed to identify associations using the rotated factor, likely reflecting the relatively limited numbers of available samples in those pathogen groups.

To interpret each MOFA factor in terms of gene expression programs, we performed GSEA on the genes ranked by their rotated MOFA factor weights for each cell type (**Fig. 3m, Supplementary Video 1**). We defined enrichment in positive weights as “positive enrichment” and enrichment in negative weights as “negative enrichment”. These analyses confirmed immunologic responses to pneumonia that have been causally implicated in pneumonia pathogenesis in murine models and humans and identified new biology^36^.

We first focused on the three MOFA factors associated with pathogen groups. Weights for “early SARS-CoV-2 factor 1” were positively enriched for genes implicated in interferon-alpha and interferon-gamma responses across all alveolar cell populations, IL6-JAK-STAT3 signaling in more mature alveolar macrophages and in CD4 and CD8 T cells, and negatively enriched for TNFα signaling via NF-κB in alveolar macrophages, CD8 T cells, and DC2 (**Fig. 3m**). Weights for “bacteria only detected factor 2” were negatively enriched for interferon response pathways across all alveolar cell populations and positively enriched for TNFα signaling via NF-κB in alveolar monocytes and monocyte-derived alveolar macrophages (**Fig. 3m**). Weights for “bacteria detected factor 3” were positively enriched for the inflammatory pathways identified in factors 1 and 2 and also were positively enriched in IL2-STAT5, complement, coagulation, allograft rejection, and immunometabolism, suggesting a synergistic effect of viral and bacterial mixed infection (**Fig. 3m**). Some pathogen-specific factors were linked to therapeutic targets. For example, genes involved in complement were positively enriched in mature alveolar macrophages in “early SARS-CoV-2 factor 1”, but negatively enriched in these cells in “bacteria only detected factor 2”. Changes in immunometabolism also demonstrated pathogen-specific differences. For example, we observed negative enrichment for genes involved in oxidative phosphorylation for all cell types and pathogens, while genes involved in glycolysis and fatty acid oxidation showed cell type- and factor-specific differences.

Weights for “immunocompromised factor 4” were negatively enriched for genes implicated in interferon responses in monocytes, alveolar macrophages, CD8 T cells, and DC2s and negatively enriched for allograft rejection across almost all cell populations (**Fig. 3c,e, Extended Data Fig. 5c, left)**. Pathways implicated in protein secretion (protein secretion, MTORC1 signaling, unfolded protein response) were positively enriched in all cell populations, particularly monocytes and alveolar macrophages, while pathways involved in oxidative phosphorylation were positively enriched in alveolar macrophages, and CD4 and CD8 T cells. These findings are consistent with the use of immunosuppressive medications that target protein translation and other immunosuppressive conditions in this population.

Male sex is associated with an increased risk of and severe outcomes from pneumonia^37^. Weights for “female sex factor 5” were negatively enriched for inflammatory signaling across almost all cell populations (TNFα signaling via NF-κB, inflammatory response, IL2 STAT5 signaling), but interferon responses were not enriched except in CD4 T cells (**Extended Data Fig. 5c, right**). Weights for “female sex factor 5” were also enriched in pathways involved in immunometabolism across cell populations. For example, oxidative phosphorylation was negatively enriched in mature alveolar macrophages and monocytes but positively enriched in CD4 and CD8 T cells. Pathways implicated in proliferation (MYC targets) and associated metabolic changes (MTORC1 signaling, unfolded protein response) were negatively enriched in monocytes and macrophages.

MOFA also provided insights into mechanisms that drive poor outcomes. For example, weights for “mortality factor 6” (**Fig. 3g**) were positively enriched for TNFα signaling via NF-κB, interferon signaling, inflammatory response, and IL6-JAK-STAT3 signaling and negatively enriched for genes involved in oxidative phosphorylation in alveolar macrophages, T cells, and DC2 (**Fig. 3m**). Weights for “ECMO factor 7” (**Fig. 3h**) were almost exclusively positively enriched for genes involved in inflammation in monocytes and monocyte-derived alveolar macrophages (TNFα signaling via NF-κB, IL2-STAT5 signaling, complement, and coagulation), and pathways implicated in cell proliferation and activation. MOFA factors were also associated with social determinants of health, including smoking status, race, and ethnicity, providing insights into their contribution to pneumonia outcomes (**Supplementary Video 1**).

We reasoned that if pathogen-specific responses in the alveolar space drive the clinical state of the patient, MOFA factors that distinguish between pathogens should also be associated with clinical features. To test this prediction, we assessed correlations between MOFA factors and continuous clinical variables extracted from the EHR (**Fig 3d, Supplementary Video 1**). “Early SARS-CoV-2 factor 1” showed a negative correlation with levels of circulating lymphocytes, confirming observational studies in patients with SARS-CoV-2 pneumonia reported early in the pandemic^23,38^. “Bacteria detected factor 3” was associated with the duration of mechanical ventilation, likely reflecting the importance of secondary bacterial infection as a driver of prolonged length of stay and outcomes in patients with SARS-CoV-2 pneumonia^38^. Finally, underscoring the association between cell type abundance and clinical parameters, the percentage of BAL fluid neutrophils was correlated with multiple factors, most notably “mortality factor 6” (**Fig. 3d**).

To further investigate the major axes of variation behind cell type-specific gene expression data, we performed PCA separately for each individual cell type. As expected, the obtained results indicated similar associations of PCA components with clinical variables as observed for the MOFA factors, with small variations among the major cell types of interest (**Extended Data Fig. 5d, e**). Both MOFA and PCA factors and their associations with clinical variables and flow cytometry-defined cell types reveal pathogen-specific host responses, suggesting pneumonia endotypes and refuting the hypothesis that pneumonia elicits a uniform immune response irrespective of the infecting organism.

### Differential gene expression analysis identifies known and novel host responses to distinct pathogens

To further explore pathogen-specific immune responses, we performed pseudobulk differential gene expression analysis within our scRNA-seq dataset. We began by identifying differentially expressed genes (DEGs) between samples from healthy controls and patients, between samples from NPCs and patients with pneumonia, and between samples with early SARS-CoV-2 infection and bacterial pneumonia (**Fig. 4a, Extended Data Fig. 6a, b, Supplemental Tables 5–34, Supplementary Video 2**). As expected, the comparisons with healthy controls revealed larger numbers of differentially expressed genes than those with NPC. For most comparisons, the largest number of DEGs was observed in alveolar macrophages (immature/maturing MRC1+C1QA–/+ and mature NUPR1+ alveolar macrophages) (**Fig. 4a**). Large numbers of DEGs between bacterial and viral pneumonia were also found in CD4 and CD8 T cells, classical monocytes-2 IL1B, and DC2. There were very few DEGs between samples coming from patients infected with gram-positive versus gram-negative bacteria or between samples from patients with *Pseudomonas* versus other gram-negative bacteria, perhaps due to patient heterogeneity and the relatively small number of samples available for individual pathogens (**Fig. 1b**, **Fig. 4a**).

**Figure 4.**
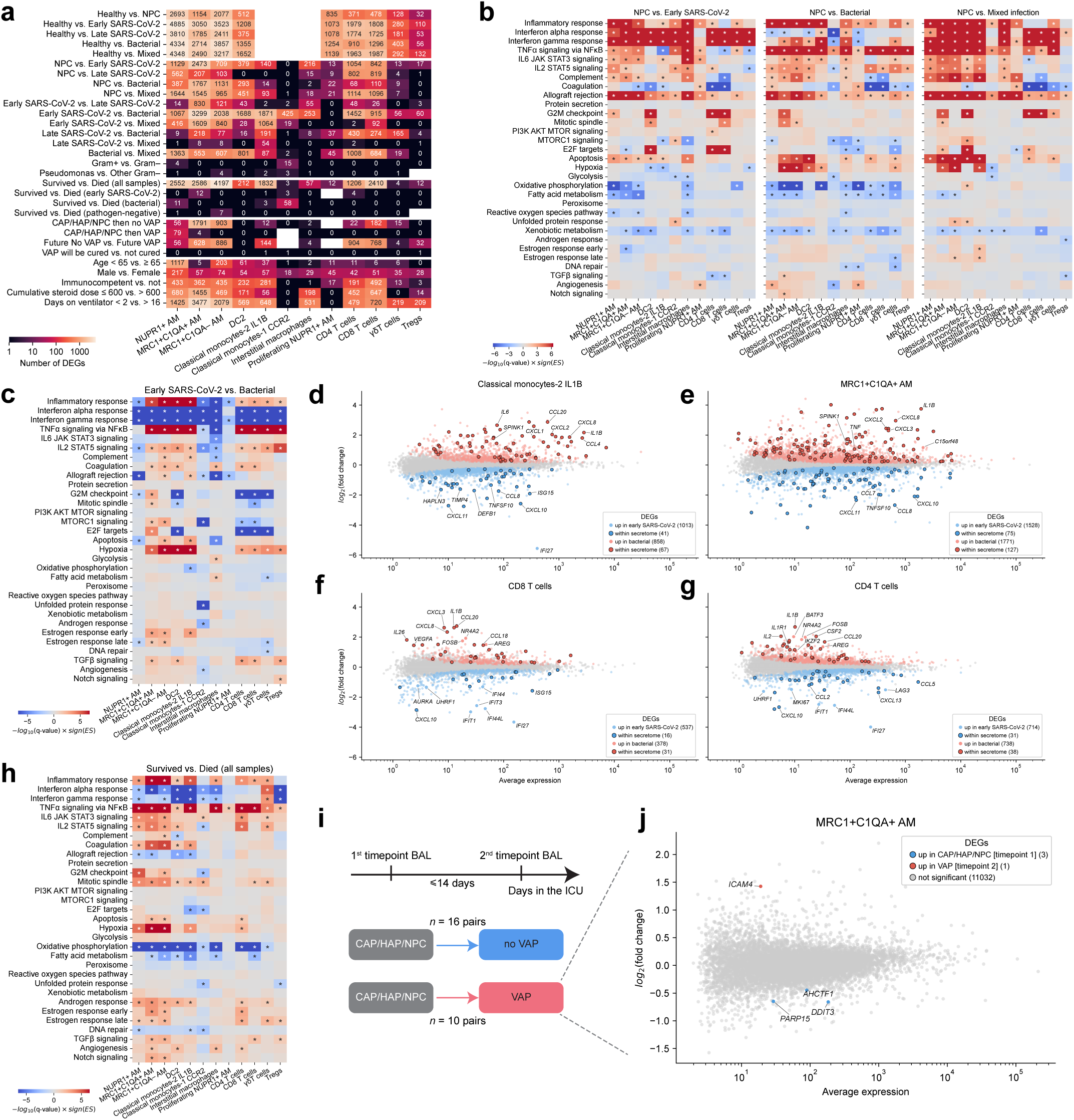
Differential gene expression identified pathogen-specific and shared responses to viral versus bacterial pneumonia. **a.** Numbers of DEGs for each comparison (rows) for more abundant cell types (columns). Empty cells indicate comparison was not performed because of the low number of samples. **b.** GSEA enrichment of hallmark gene sets (rows) in gene lists ranked by log-fold change per each cell-type (columns) for comparisons of sample groups “NPC” vs “Early-SARS-CoV-2, “NPC” vs “Bacterial”, and “NPC vs Mixed”. * indicates significant enrichment using GSEA after FDR correction. **c.** GSEA enrichment of hallmark gene sets (rows) in gene lists ranked by log-fold change per each cell-type (columns) for comparisons of sample groups “Early SARS-CoV-2 vs Bacterial” * indicates significant enrichment using GSEA after FDR correction. **d–g.** MA plots of differential gene expression analysis between bacterial and early SARS-CoV-2 samples for Classical monocytes-2 IL1B (**d**), MRC1+C1QA+ AM (**e**), CD8 T cells (**f**) and CD4 T cells (**g**). Red circles: genes with significantly increased expression in bacterial infection; blue circles: significantly increased expression in early SARS-CoV-2 samples; black circles: genes that encode proteins annotated as secreted in the human protein atlas. Significance determined by DESeq2 test with FDR correction. **h.** GSEA enrichment of hallmark gene sets (rows) in gene lists ranked by log-fold change per each cell-type (columns) for comparisons of sample groups “Survived” vs. “died”. * indicates significant enrichment using GSEA after FDR correction. **i.** Schematic describing serial sample analysis. Serial samples were distinguished by whether a second sample collected after the first was adjudicated as a VAP or not a VAP irrespective of whether the initial paired sample was adjudicated as community acquired pneumonia (CAP), hospital acquired pneumonia (HAP) or NPC. **j.** MA plot of differential gene expression analysis when comparing serial samples from patients adjudicated as VAP to the paired initial sample. Red circles indicate significant increased expression in VAP at timepoint 2, and blue circles indicate significant increased expression at the paired initial sample. Significance determined by DESeq2 test with FDR correction.

We performed GSEA on the lists of genes ranked by log-fold change to determine significant associations with specific pathways (**Fig. 4b**). Pathways involved in interferon-alpha and -gamma responses were enriched in early SARS-CoV-2 pneumonia compared with NPC across multiple cell populations, and IL6-JAK-STAT3 signaling and IL2-STAT5 signaling were enriched in alveolar macrophages and monocytes (**Fig. 4b**). These findings parallel those obtained from scRNA-seq of nasal samples from healthy people after experimental SARS-CoV-2 infection^39^. Similar findings were observed between samples from patients with early SARS-CoV-2 pneumonia and healthy controls (**Extended Data Fig. 6c**).

In contrast, samples from patients with bacterial pneumonia showed reduced expression of genes contributing to pathways involved in interferon alpha responses in alveolar macrophages relative to NPC (**Fig. 4b**). Surprisingly, these pathways were reduced even in comparison to healthy controls, consistent with downregulation of interferon responses by NFκB activation (**Extended Data Fig. 6c**)^40^. Genes involved in the TNF signaling via NFκB pathway were upregulated in all cell populations relative to NPC (**Fig. 4b**). Similarly, pathways implicated in inflammatory response, IL2-STAT5 signaling, and allograft rejection were upregulated in bacterial pneumonia when compared with NPC in most cell populations (**Fig. 4b**). These pathways were not upregulated in classical monocytes-1 CCR2, which are sampled in peripheral blood. Distinct from samples from patients with early SARS-CoV-2 pneumonia, changes in pathways related to proliferation and immunometabolism in T cells were limited in patients with bacterial pneumonia (**Fig. 4b**). We observed a similar pattern of changes when samples from patients with bacterial pneumonia were compared with healthy controls (**Extended Data Fig. 6c**).

When compared with NPC, samples from patients with mixed viral and bacterial pneumonia showed upregulation of a summation of pathways associated with SARS-CoV-2 and bacterial pneumonia (**Fig. 4b**). Interferon-alpha and -gamma responses, as well as TNFα signaling via NF-κB, were upregulated across alveolar macrophage and T cell populations. Mixed infection was also associated with activation of pathways involved in complement. The oxidative phosphorylation pathway, which was downregulated in patients with viral or bacterial pneumonia, was unchanged in most cell types in patients with mixed pneumonia (**Fig. 4b**). Despite the small number of DEGs between gram-positive and gram-negative bacteria, GSEA identified distinct signaling pathways across alveolar cell populations that suggest possible pathogen-specific responses (**Extended Data Fig. 6d**).

We next sought to determine whether our analysis identified biologically meaningful differences at the level of individual genes. Because most patients with pneumonia do not require mechanical ventilation, blood-based biomarkers of the host response are preferred to those based on BAL sampling. Accordingly, we focused on genes encoding secreted proteins, as defined in the Human Protein Atlas (**Supplementary Table 35**), that were differentially expressed in viral compared with bacterial infection in alveolar macrophages or T cells. For example, MRC1+C1QA+ AMs from early SARS-CoV-2 samples showed increased expression of *CXCL11*, *CXCL10*, *CCL8*, *CCL7 and TNFSF10*, while bacterial infection was associated with increased expression of *TNF*, *IL1B, CXCL8, CXCL2, CXCL3, SPINK1,* and *C15orf48* (**Fig. 4c–g, Supplementary Table 15**). Many of these genes have been identified in other studies^41–43^ but others represent novel potential biomarkers. Many of the observed DEGs are chemokines or cytokines, which are involved in the recruitment of specific inflammatory cell populations into the alveolar space^44^. This finding may explain the association of multiple MOFA factors with BAL fluid cell type abundance measured by flow cytometry and scRNA-seq (**Fig. 3c**).

Investigators recently used Olink technology to generate a comprehensive proteomic analysis of blood samples from patients with 59 distinct diseases, including influenza A infection, allowing us to credential our findings in an independent dataset of blood measurements^45^. They found that CCL8, RIGI (*DDX58* in our dataset, originally annotated as not secreted in the Human Protein Atlas), and LTA4H best distinguished patients with influenza infection from those with other infections. In our data, *CCL8* is expressed in Classical monocytes-2 IL1B, MRC1+C1QA– and MRC1+C1QA+ AMs, where it is upregulated in early SARS-CoV-2 pneumonia compared with bacterial pneumonia (**Fig. 4c–f, Supplementary Video 2**). RIGI (*DDX58*) is widely expressed in our dataset and is upregulated in monocyte-derived alveolar macrophages and CD4 and CD8 T cells in patients with early SARS-CoV-2 pneumonia compared with bacterial pneumonia (**Fig. 4c–f, Supplementary Video 2**). LTAH4 is responsible for production of the potent neutrophil chemoattractant LTB4, is expressed in monocytes and alveolar macrophages in our dataset and is upregulated in patients with bacterial pneumonia (**Fig. 4c–g, Supplementary Video 2**).

We performed additional comparisons to gain insights into the biological mechanisms underlying clinical features associated with pneumonia outcomes. In samples from patients ≥ 65 years of age compared with those < 65 years of age, we detected a large number of DEGs in alveolar macrophages (**Fig. 4a**) but few age-related DEGs in CD4 or CD8 T cells (**Fig. 4a**). Age ≥ 65 years was associated with increased interferon signaling in circulating monocytes and non-classical monocytes but reduced interferon signaling in MRC1+C1QA+ and MRC1+C1QA– AMs and monocyte-derived alveolar macrophages (**Extended Data Fig. 6e**). Immunocompromise was associated with reduced inflammatory signaling, including interferon responses and TNFα signaling via NF-κB in monocyte-derived alveolar macrophages, but greater TNFα signaling via NF-κB, IL-6 JAK STAT3 signaling, and IL2 STAT5 signaling in NUPR1+ AMs and CD4 T cells (**Extended Data Fig. 6f**). Treatment with lower doses of corticosteroids was associated with higher interferon responses across most cell types (**Extended Data Fig. 6g**).

We then examined genes that were differentially expressed in BAL samples from patients who subsequently lived compared with those who died in the hospital. When including all samples, the number of differentially expressed genes was large, particularly in alveolar macrophages, T cells, and recruited classical monocytes (**Fig. 4a**). Across all these populations, GSEA revealed significant upregulation of TNFα signaling via NF-κB and reduced interferon-alpha and interferon-gamma responses (**Fig. 4h**). We repeated the same analysis in patients with bacterial pathogens, viral pathogens, and pathogen-negative pneumonia. While the number of differentially expressed genes was much lower (nearly absent in bacterial pneumonia), GSEA identified reduced interferon alpha response in all alveolar macrophage populations and increased TNFα signaling via NF-κB in alveolar macrophages and CD4 T cells from patients who died compared with those who survived, irrespective of pathogen group (**Supplementary Video 2**). These data suggest that a persistent hyperinflammatory state in the alveolus, likely driven by bacterial infection or superinfection, is associated with poor outcomes.

We applied Gene Set Variation Analysis (GSVA)^19^ to enable a more detailed view into pathway enrichment in gene expression profiles within individual samples (**Supplementary Video 3**).

### Serial BAL sampling identifies robust immune responses to secondary bacterial pneumonia

Patients with pneumonia complicated by sepsis have been reported to develop suppression of innate and adaptive immune systems to varying degrees^46,47^. A prediction from this hypothesis, sometimes referred to as “immunoparalysis”, is that patients with severe pneumonia receiving mechanical ventilation will have a dampened immune response during a subsequent VAP episode when compared with their previous pneumonia episode. Accordingly, we identified paired samples from patients with or without pneumonia who underwent a subsequent BAL sample for suspected VAP. We then compared these subsequent BAL samples to the first sample according to whether the subsequent sample was adjudicated to define a VAP or not (n = 10 CAP/HAP/NPC then VAP, n = 16 CAP/HAP/NPC then no VAP, **Fig. 4a,i**). Patients who did not have VAP diagnosed on their subsequent BAL sample exhibited a robust DEG signature compared with their first BAL sample in alveolar macrophages, specifically immature alveolar macrophages, and CD8 T cells when comparing their initial versus subsequent BAL sample, suggesting resolution of alveolar inflammation. In contrast, patients who had VAP diagnosed on their subsequent BAL sample had a gene expression profile resembling their initial BAL, indicating a similar response to their initial and subsequent pneumonia (**Fig. 4j**). This result argues against the immunoparalysis hypothesis, which predicts that the immune response to a subsequent pneumonia would be impaired, resulting in differential gene expression.

We also compared the initial BAL samples in patients who went on to develop VAP (Future VAP) or not (No Future VAP). These samples differed in their gene expression profiles, potentially indicating signatures that predict or predispose to VAP onset. Surprisingly, interferon responses were upregulated in patients who did not develop a subsequent VAP, possibly due to more patients with early SARS-CoV-2 pneumonia in the subset of patients who did not develop VAP versus those who did (9 versus 3, respectively) (**Supplementary Video 2**).

### Host response on the single cell level classifies pneumonia etiology

To determine whether the single-cell transcriptome of individual immune cells in BAL fluid reflects pathogen-specific responses, we employed Geneformer^17^. Single cell embeddings of pretrained Geneformer clustered by cell types (**Fig. 5a**). Embeddings for cells coming from distinct pathogen-infected samples occupied different regions of the latent space (**Fig. 5a–b**).

**Figure 5.**
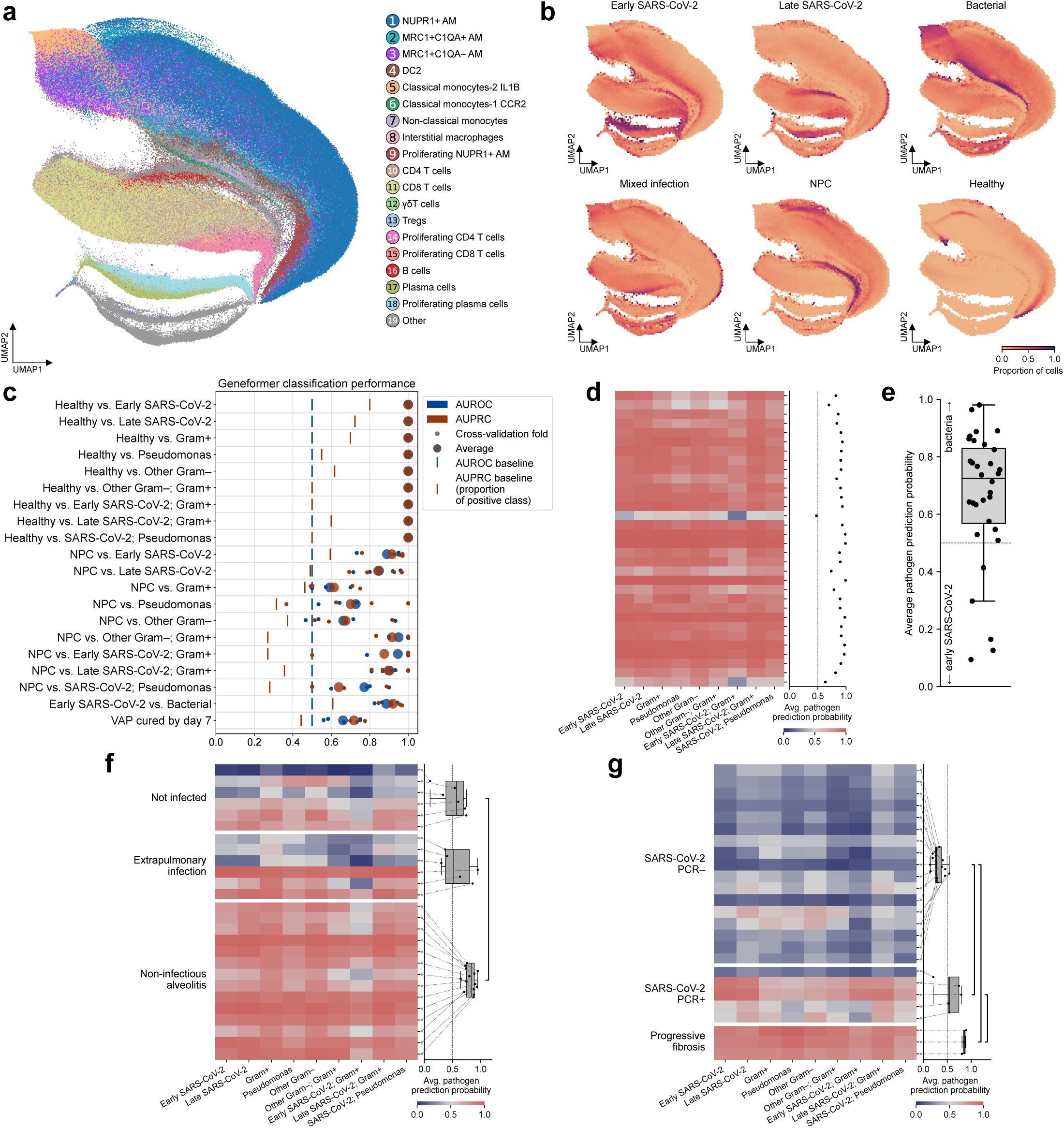
Pathogen-specific host response at the single-cell level using a foundation model. **a.** UMAP of pretrained Geneformer embeddings of single cells, colored by cell types within the scRNA-seq data. **b.** Hexbin UMAP of pretrained Geneformer single-cell embeddings, with each hexagon colored by the proportion of cells from the indicated pathogen group. **c.** Classification performance of fine-tuned Geneformer models evaluated by aggregating cell-level predictions at the sample level. For each classification task (y-axis) the area under the receiver operating characteristic curve (AUROC), area under the precision recall curve (AUPRC) is presented. For each of the 4 cross-validation folds, the classification performance was averaged over 5 random initializations. Baseline values for AUROC (0.5) and AUPRC based on class imbalance are shown. **d.** Sample-level classification probabilities from fine-tuned Geneformer models for the 32 samples (rows) that were clinically adjudicated as pneumonia in the absence of a detected pathogen. The heatmap (left) shows sample level prediction for each individual pathogen (columns). The average probability for classification as any pathogen is shown on the right. **e.** Sample-level classification probabilities from fine-tuned Geneformer models distinguishing bacterial pneumonia from early SARS-CoV-2 infection across the 32 samples that were clinically adjudicated as pneumonia in the absence of a detected pathogen. **f.** Sample-level classification probabilities from fine-tuned Geneformer models for the 26 non-pneumonia control (NPC) samples. NPC episodes were further adjudicated as having respiratory failure from a non-infectious etiology (not infected), an extrapulmonary infection, or a non-infectious etiology. (Left) Heatmap of sample level prediction for each individual pathogen. (Right) Box-and-whiskers plots of the average probability for any pathogen for each of the three subclassifications of NPC. Bracket denotes statistically-significant differences (*q*-value < 0.05, Mann-Whitney U tests with FDR correction). **g.** Fine-tuned Geneformer pathogen classification of previously published scRNA-seq data from BAL fluid. Data were derived from 25 BAL samples collected from patients with persistent respiratory symptoms and chest computed tomography (CT) abnormalities following clinical recovery from SARS-CoV-2 pneumonia. Samples are grouped by SARS-CoV-2 PCR results on BAL fluid. Three samples from two patients who subsequently required lung transplantation for progressive post-SARS-CoV-2 pneumonia pulmonary fibrosis are shown as a separate group. (Left) Heatmap of fine-tuned Geneformer predicted probabilities for individual pathogens at the sample level. (Right) Box-and-whisker plots of the mean predicted probability of detecting any pathogen across groups. Brackets denote statistically-significant differences (*q*-value < 0.05, two-sided Mann-Whitney U tests with FDR correction).

We fine-tuned Geneformer to classify individual cell types as coming from samples infected by one of the 9 pathogen groups, and to recognize viral versus bacterial infection. In addition, we fine-tuned Geneformer for cell-level prediction of VAP cure within 7 days. With AUROC close to 1, individual cell populations showed high predictive value for distinguishing healthy controls from patients with pneumonia (**Extended Data Fig. 7a**). The AUROC values were highest for MRC1+C1QA+ and MRC1+C1QA– AMs and classical monocytes-2 IL1B, reflecting the biological role of macrophages as sensors of their microenvironment^12,48^. Among the classification tasks with NPC, NPC versus early SARS-CoV-2 pneumonia also obtained high AUROC scores, which were the highest (>0.8) for MRC1+C1QA+ AM, CD4 T cells, Tregs and DC2 (**Extended Data Fig. 7b**). In classification of viral versus bacterial pneumonias, the model obtained high average AUROC scores for the majority of cell types, especially alveolar macrophages, with a lower performance for the proliferating T cells. Performance in the VAP cure prediction task had the highest AUROC for proliferating CD8 T cells, followed by Tregs (**Extended Data Fig. 7c**).

We next averaged cell-level predictions for each sample and assessed classification performance on a per sample level. The sample-level classifiers obtained good results, with AUROC and AUPRC close to 1 on the test set for the tasks of distinguishing pathogen-infected samples from healthy controls (**Fig. 5c, Supplementary Table 36**). Similar to single cell-level classifiers, the performance decreased for the tasks of distinguishing samples from patients with infection from NPC, but still remained high compared with random baselines. With AUPRC of 0.72 and AUROC 0.66, Geneformer performance for VAP cure prediction exceeded the random baseline.

Twenty-two samples in our dataset were clinically adjudicated as having pneumonia caused by known pathogens that were not included in our primary analysis (**Fig. 1b**). Our models accurately classified 95% of these samples as pneumonia (defined as average predicted probability > 0.5 across pathogen models), and accurately identified the pathogen type (viral or bacterial) in 78% of the samples (7 out of 9, **Extended Data Fig. 7d, e**). Another set of 32 samples was clinically adjudicated as pneumonia by the expert panel but no pathogen was identified. Our models predicted that the majority (96%) of these samples were infected, with 85% predicted as coming from patients with bacterial pneumonia (**Fig. 5d,e**).

Samples adjudicated as NPC include patients with disorders that mimic pneumonia. The resulting alveolar inflammation might explain the reduced performance of our models in distinguishing NPC from pneumonia compared with healthy controls. To test this hypothesis, we used our clinical adjudication to subcategorize NPC samples as those with a non-infectious inflammatory condition (non-infectious alveolitis), those with an identified extrapulmonary infection, and those with a non-inflammatory cause of respiratory failure (not infected, e.g., heart failure or atelectasis) (**Extended Data Table 5**). When we applied the healthy versus pathogen group classifiers to the NPC samples, 17/20 samples from patients with an extrapulmonary infection or a systemic inflammatory disorder were classified as not healthy by our models, and the average pathogen prediction probability for samples with alveolitis was higher than that for samples with respiratory failure from a non-inflammatory cause (**Fig. 5f**).

We wondered whether our models would make reasonable predictions when applied to BAL fluid data from patients who were neither healthy nor critically ill. Accordingly, we tested the performance of our models on previously published scRNA-seq data generated from BAL samples obtained from patients with persistent respiratory symptoms and CT abnormalities more than 3 months after COVID-19, a subset of patients with prolonged acute sequelae of COVID-19 (PASC or “long COVID”)^12^. Our models classified 68% of these samples as negative for pneumonia (17/25, **Fig. 5g**). In the samples from these patients in which SARS-CoV-2 was detected by nasal swab PCR, our model identified 4 out of 5 as infected. In the three samples from the two patients who progressed to require lung transplantation for COVID-19, our models predicted infection with high probability. Overall, this performance across distinct datasets demonstrates that single-cell transcriptomes contain pathogen-specific signatures, supporting the conclusion that these signatures represent pneumonia endotypes.

## Discussion

Using deep molecular profiling of bronchoalveolar lavage fluid from a large, well-phenotyped cohort of mechanically ventilated patients, we showed that severe pneumonia is characterized by distinct, pathogen-specific host response endotypes within the human lung. These endotypes were evident across multiple modalities, including alveolar immune cell composition, cell-type–specific transcriptional programs, and routinely collected clinical features available in the electronic health record, and they evolved dynamically over the course of critical illness. Pathogen-associated differences in interferon and NFκB signaling, inflammatory and metabolic pathways, and immune cell recruitment, particularly among monocytes, alveolar macrophages, neutrophils, and T cell subsets, were consistently linked to clinical trajectories and outcomes, such as ventilator-associated pneumonia, organ failure, and mortality, even among immunocompromised patients. Our results suggest that endotypes defined by host-pathogen interactions could inform the clinical management of severe pneumonia rather than relying on syndromic definitions.

Pneumonia is the second most common cause of sepsis syndrome, defined as life-threatening organ dysfunction caused by a maladaptive host response to infection, and is the most common cause of death from sepsis^49,50^. Investigators have suggested that the multiple organ dysfunction that develops in patients with sepsis results from immunoparalysis, where immune cell exhaustion leads to secondary infections and a loss of mucosal barrier function^46,47^. Irrespective of pathogen, we observed evidence of T cell and alveolar macrophage activation in patients who died from pneumonia compared to those who survived, failing to detect evidence of immunoparalysis. In serial samples, we observed preserved rather than diminished inflammatory signaling in response to a second pneumonia episode during the same course of critical illness. These data suggest that, in the majority of patients, persistent immune activation rather than immunoparalysis is associated with poor outcomes. Our data point to specific immunologic pathways associated with persistent immune activation and adverse outcomes. Prominent among these are processes that regulate metabolism and stress responses in alveolar macrophages and T cells that can be causally tested in murine models of pneumonia.

Our results extend the work of Calfee and colleagues, who used latent class analysis to identify “hyperinflammatory and hypoinflammatory phenotypes” in patients with ARDS or sepsis^51^. Patients with a hyperinflammatory phenotype had a higher mortality and were more likely to respond to interventions in post-hoc analyses of interventional clinical trials. However, this classification relies on variables not routinely measured in clinical practice and does not include immunocompromised patients, an important ICU patient population who suffer from disproportionate mortality and are frequently excluded from clinical trials. We provide strong evidence that in pneumonia, this hyperinflammatory phenotype is driven by exuberant alveolar inflammation, often associated with bacterial infection. Furthermore, our molecular profiling includes immunocompromised patients, providing foundational data for developing next-generation, blood-based classifiers that include this important ICU population.

Our study has several limitations. First, while ours is the most comprehensive dataset phenotyping the clinical, cellular, and molecular features of the alveolar space in patients with pneumonia available to date, the size of our single-center cohort provided limited statistical power to detect differences in transcriptional signatures between patients with specific pneumonia pathogens, for example *P. aeruginosa* compared with other bacterial infections. Secondly, the BAL procedure does not sample parenchymal cells in the alveolar space that play important roles in pneumonia pathogenesis^52^. Emerging spatial technologies applied to autopsy or lung biopsy tissues will provide important complementary information to our study. Third, the observational design of our study results in relative oversampling of patients with prolonged courses of mechanical ventilation who are at increased risk of VAP and poor outcomes, highlighting the need for studies that sample patients with resolving pneumonia. Fourth, we used state-of-the art tools to diagnose pneumonia, adjudicate outcomes, and identify the causative pathogen in our cohort. Nevertheless, in a substantial fraction (31%) of our cohort, the pneumonia pathogen was not identified. More sensitive tests, for example cell free microbial DNA analysis, might address this limitation^53^. Fifth, while we did not detect a signal of immunoparalysis in serial samples, the number of serial samples was small and the infecting pathogens varied. Larger studies using serial samples can address this limitation. Finally, current foundation models like Geneformer show promise as classification tools but have limited interpretability. New data science approaches are needed to leverage these tools for therapeutic discovery. To that end, we have made our data publicly available for secondary analysis as newer models and data science approaches emerge.

In conclusion, our findings suggest that a molecular approach to diagnosis and therapy could guide the selection and timing of antimicrobial and immunomodulatory therapies to improve outcomes for patients with severe pneumonia. By limiting antibiotic overuse and identifying patients who may benefit from host-directed therapies, this approach also addresses the growing public health challenge of antimicrobial resistance and unclear indications for immunomodulatory agents. The identification of specific therapies to treat patients with a molecular diagnosis of SARS CoV-2 pneumonia during the COVID-19 pandemic demonstrated the value of this paradigm^8,54–60^. Our results further suggest that small biomarker-guided trials enabled by rapid pathogen and host-response diagnostics may outperform large trials based on syndromic definitions by increasing the likelihood of endotype-specific benefit while minimizing harm induced by overtreatment.

## Methods

### Cohort

We leveraged data and samples from the Successful Clinical Response in Pneumonia Therapy (SCRIPT) study. SCRIPT is an NIH (NIAID)-funded prospective single-center observational cohort study that models pneumonia pathobiology using distal lung samples from patients with severe pneumonia at Northwestern Memorial Hospital. The study began enrolling patients in June 2018 and was supported by the Northwestern University Trustees during the freeze in NIH funding to Northwestern University between March 26, 2025 and December 10, 2025.

Patients were eligible for enrollment in the SCRIPT study if they were critically ill, receiving mechanical ventilation, underwent BAL for suspected pneumonia, and were followed through to hospital discharge. The patient’s family or legal authorized representative provided informed consent. This study is approved by the Northwestern IRB under Study ID STU00204868. More details of the SCRIPT study are available in previous manuscripts.^26,38^ Healthy volunteers were enrolled in studies STU00206783 and STU00214826 at Northwestern University, Pro00088966 and Pro00100375 at Duke University.

Single-cell RNA-sequencing data included BAL samples from the SCRIPT cohort as well as additional BAL samples from external cohorts: 9 samples from healthy individuals, 25 samples from 24 patients with post-acute sequelae of COVID-19^12^, and 4 BAL samples from 2 patients enrolled in a separate clinical trial of a CRAC channel inhibitor Auxora (IRB STU00214054).

A small subset of patients had multiple enrollments; each new enrollment was given a new study ID. Eleven patients who underwent lung transplantation during the index hospitalization were adjudicated as having died on the day of the transplantation, and days after lung transplantation were censored.

### Clinical data

For the SCRIPT cohort, we obtained electronic health record (EHR) data via the Northwestern Enterprise Data Warehouse^61^ and summarized it on a daily basis for 15,287 ICU-patient-days as previously described^38,62^. Features were selected by pulmonary and critical care physicians to reflect parameters commonly reviewed in daily ICU rounds, including vitals, ventilator settings, laboratory results, and organ support devices^22^. We also provided summative information such as days of mechanical ventilation, cumulative antibiotic spectrum exposure, and cumulative steroid exposure. Additional details were captured by the study research team by interviewing the patient’s family or abstracting from the EHR through standardized forms in REDCap. Mortality was defined as patients who died in the hospital, discharged to hospice, or underwent lung transplantation for refractory respiratory failure during the index hospitalization.

Pneumonia episode adjudications were performed by a panel of pulmonary and critical care attending physicians using a published procedure^13^. From these adjudications, we extracted pneumonia episode start date and the corresponding BAL identifier, episode etiology, category (CAP, HAP, VAP), duration (if available) and whether the episode was cured, not cured or indeterminate.

### BAL sample collection and use in the ICU

Testing on BAL fluid is clinician-directed and guided by clinical protocols and order sets in our ICU that include automated cell counts, biochemical analyses, PCR for common pneumonia pathogens and selected antimicrobial resistance genes, and quantitative microbial cultures. These studies are complemented by blood and urinary tests to detect specific pathogens (e.g., serum levels of (1-3)-β-D-glucan and urinary antigens for *Streptococcus pneumoniae* and *Legionella pneumophila*). We have shown that our ICU clinicians adjust antimicrobial therapy based on BAL fluid and urinary test results to enhance antibiotic stewardship and reduce exposure to broad-spectrum antibiotics compared with guideline-based empirical therapy^63,64^.

Bronchoscopic and non-bronchoscopic BAL samples^25^ were processed for pathogen testing (quantitative bacterial culture, fungal culture, susceptibility testing, multiplex PCR with the Biofire Pneumonia Panel^65^ or Respiratory Pathogen Panel) and clinical cell type differential. Details of procedures regarding sample collection and processing were as previously described.^25,26,38^

### Sample labels for machine learning

To define infecting lung pathogen labels, we summarized pathogen testing results recorded in the EHR for each BAL sample, including quantitative bacterial culture, susceptibility and PCR testing, and the percentage of neutrophils in the clinical cell count differential using our previously published adjudication protocol^13^. A BAL sample was considered “pathogen identified” if a pathogen was detected by PCR or in bacterial culture above 1000 CFU, or at any CFU level when the sample contained ≥ 50% neutrophils. For some pathogens, detection at any CFU level was considered positive (**Supplementary Table 37**). Based on the set of all pathogens detected in a sample we derived hierarchical pathogen group labels (visualized in **Fig. 1b**). We distinguished “viral” infections, consisting of infections with SARS-CoV-2 in the absence of bacterial co-infection (60 scRNA-seq samples), viral and bacterial coinfections (“mixed”, 33 scRNA-seq samples), and “bacterial” infections in the absence of virus (58 scRNA-seq samples).

From this hierarchy, we selected groups (at various levels) represented by at least eight samples in the scRNA-seq subset of the data to ensure sufficient statistical power for downstream analyses. Within the group of samples from patients with SARS-CoV-2, we compared early (within 48 hours of intubation) with late (>48 hours of intubation) infection, aligning with the syndromic definition of ventilator-associated pneumonia (VAP)^4^. Gram-positive and gram-negative bacteria were reported to induce distinct transcriptional signatures in circulating inflammatory cells based on constituents of their cell wall^66^. *Pseudomonas spp.* (here simply referred to as *Pseudomonas*) are common pathogens implicated in VAP^4^. This resulted in 9 defined pathogen groups: “Early SARS-CoV-2”, “Late SARS-CoV-2”, “Gram+”, “*Pseudomonas*”, “Other Gram–” (excluding *Pseudomonas*), “Other Gram–; Gram+”, “Early SARS-CoV-2; Gram+”, “late SARS-CoV-2; Gram+”, and “SARS-CoV-2; *Pseudomonas*”.

22 scRNA-seq samples with “rare pathogens”, and combinations of rare pathogens, identified in a small number of samples, were excluded from pathogen-based analysis Out of the 40 scRNA-seq samples obtained from patients with pneumonia that did not contain identified pathogen (“no pathogen identified”), we distinguished 32 from patients with pneumonia lacking an identified pathogen (“pathogen-negative” pneumonia) and 8 from patients with pneumonia who had a previous sample containing a pathogen (“pathogen cleared”). Of the 26 scRNA-seq samples adjudicated as “NPC”, 6 were from patients with a known or suspected infection outside the lung (extrapulmonary infection), 14 were from patients with a non-infectious condition known to induce a systemic inflammatory response (non-infectious alveolitis), and 6 were from patients with causes of respiratory failure that are not associated with high-grade systemic inflammation (not infected, e.g., heart failure or atelectasis).

Additionally, we considered other pathogen information for samples (not visualized in **Fig. 1b** for clarity): samples that contained a bacterial pathogen as detected by PCR or culture (“bacteria detected”, 113 scRNA-seq samples), contained bacteria as the sole pathogen(s) detected (“bacteria only detected”, 66 scRNA-seq samples), contained viral pathogen as detected by PCR (“virus detected”, 119 scRNA-seq samples), contained virus as the sole pathogen(s) detected (“virus only detected”, 72 samples). Clinical adjudication protocol^13^ also defined pneumonia episode pathogen etiology (“adjudicated: viral”, 52 scRNA-seq samples, “adjudicated: bacterial”, 43 scRNA-seq samples, “adjudicated: mixed”, 47 scRNA-seq samples, “adjudicated: negative” [20 scRNA-seq samples; pneumonia in the absence of identified pathogen]), which took into account other information from the EHR and clinical notes, besides BAL pathogen results, but was available only for episode-defining BALs.

We used the BAL-based pathogen identification and categorization to define infecting lung pathogens on the days between the BALs as having the same pathogens as at the last BAL (i.e., assuming that pathogens remain the same between consecutive BALs). Pathogen labels were set to NA for patient-days before the first BAL, and after ICU readmission until first BAL during readmission. We set patient-days after the patient adjudicated pneumonia cure to no pathogens. Additionally, we labeled BALs from adjudicated pneumonia episodes, in which pathogen testing was negative and was not preceded by a pathogen positive BAL in that episode as “pathogen-negative” BALs.

We formulated the VAP onset prediction task as predicting VAP status of the next BAL within 7 days, i.e., whether the next BAL will detect a new bacterial VAP episode, as adjudicated by our expert panel. Positive labels (‘’VAP onset within 7 days’’) were assigned to 1566 patient-days –7 to –2 before a bacterial VAP BAL sample initiating a new VAP episode (under the assumption that patients on day –1 before a VAP are highly likely to already have VAP) or a BAL sample with “superinfection” adjudication from a CAP or HAP episode (which were set only when a repeat BAL was performed). Negative labels (“no VAP onset within 7 days”) we assigned to 251 patient-days –7 to –1 preceding an NPC BAL sample or a BAL sample adjudicated as “cure”, “prophylaxis” or “cure + prophylaxis” during CAP or HAP episodes. All other patient-days received NA labels. When both labels were applied to the same patient-day, the “no VAP onset within 7 days” label took precedence. For example, if patient had a BAL on day 3 that was adjudicated as NPC, and a BAL on day 6 that was adjudicated as VAP, day 1 would qualify for both labels; however, the earlier NPC BAL rules out VAP onset within the subsequent 7 days, and the day was therefore labeled as “no VAP onset within 7 days”.

“VAP cure” status labels were defined for the first 3 days of each VAP episode and indicated whether the episode was adjudicated as cured at the day 7 assessment (297 patient days) or not (335 patient days). Episodes with indeterminate day 7 adjudication received NA labels. “VAP cure” labels for scRNA-seq BAL samples were taken from their corresponding patient-days (19 VAP cured, 22 VAP not cured).

Patient-day and sample counts for all labels defined above are summarized in **Supplementary Table 1**.

### Data split for model training

We used the three labels defined above to perform supervised machine learning. To assess classification performance of our models, we split our data into training and test sets using 4-fold cross-validation. Splitting was performed at patient level, such that all patient-days for each patient were assigned entirely in the train or in the test portion of the fold. For each fold and each label value, approximately 22% of patients with that value were randomly assigned to the test set. We first generated the splits within the scRNA-seq subset of the cohort and then extended these patient-level assignments to the full cohort. The exact fraction of patients in the test set varied (14–29% in some combinations of label, value, and fold), as some patients had multiple values for the same label throughout their ICU stay and due to the two-step splitting procedure.

### EHRformer

We used EHRformer^14^, a transformer-based model for vectors of clinical variables. As the base architecture for EHRformer we set the standard Transformer Encoder^67^ with 4 transformer layers, embedding dimension of 64, intermediate layer size of 128, and 4 attention heads^14^.

### Input representation

To represent data for a single patient-day, we binned numerical values into 4 bins using quartile binning. For features with few unique values (e.g. Glasgow Coma Scale verbal response score or number of PEEP changes in a day) custom binning was used instead. Binary categorical features (e.g. intubation flag, sex) were assigned to extreme bins. If a feature value was absent for the patient-day, we assigned a dedicated NA bin for it. Each EHR feature and each of the five possible bins (four value bins plus NA) were associated with learnable vector embeddings. The representation of a feature value was defined as the element-wise sum of the corresponding feature embedding and bin embedding. Additionally, we used two special tokens with learnable embeddings: MASK token, which replaced the bin token for the masked features during pretraining (see below), and CLS token, which we prepended for each patient-day sequence. In total, each patient-day was represented with a sequence of 51 tokens: CLS and 50 tokens for each EHR feature, encoding both the feature identity and its binned value.

### EHRformer pretraining

Following the BERT pretraining procedure^67^, we pre-trained EHRformer using a masked language modeling objective. During this phase, we randomly masked 15% of the values of each EHR entry by replacing the true bin embedding with the learnable MASK embedding. We did not mask CLS token or NA bin token. We trained the model to reconstruct the masked tokens for 300 epochs using the AdamW optimizer with a linear learning rate schedule. To prevent test data leakage, we pretrained separate models for each combination of modeling label and cross-validation fold (see “Data split for model training” above and “EHRformer finetuning and evaluation” below), resulting in 12 pretrained models (3 labels × 4 folds). Additionally, we pretrained one model on all data to be used in perturbation analysis (see “EHRformer perturbation analysis” below).

### EHRformer perturbation analysis

To analyze the relationships between clinical features that the model learned during pretraining only, we performed a perturbation analysis, in which model predictions on perturbed inputs were compared with predictions on the original inputs. We used the EHRformer model pretrained on the whole dataset and ran a forward pass on all data to obtain unperturbed predictions for each patient-day and each clinical feature, corresponding to predicted quartile bins. We defined a perturbation by the clinical feature and a direction of change, e.g. increasing PEEP. Following this example, we split the data into 3 groups based on the quartiles of the PEEP feature, excluding those already in the maximum quartile. Within each group we set the PEEP feature to the maximum quartile and ran the forward pass on this perturbed data to obtain predictions for quartiles of the remaining clinical features. For each patient-day we subtracted the unperturbed predicted quartile from the perturbed predicted quartile and normalized it to the difference between the source and target quartiles for the perturbed feature. These normalized differences were averaged within each source quartile. We excluded patient-days for which the model did not predict one of the quartile bins for perturbed or unperturbed data.

### EHRformer feature analysis

To analyze the learned latent space of EHRformer solely after pretraining, we extracted patient-day embeddings from one of the pretrained models (pathogen group label, cross-validation fold 3). We ran a forward pass on the full dataset using that model, and computed the average embedding of all 50 EHR features from the penultimate transformer layer of the model (excluding the CLS token). In this way we obtained 64-dimensional embeddings for every patient-day in the EHR data. We then projected the embeddings to 2 dimensions using UMAP with min_dist=0.1, spread=0.5, n_neighbors=8.The resulting reduced embedding space was visualized using hexbin density plots using matplotlib to truthfully represent regions with high densities of closely overlapping data points.

### EHRformer finetuning and evaluation

We finetuned a total of 240 EHRformer models to evaluate performance and stability. Each evaluation was formulated as a binary classification task: 9 NPC vs. a specific pathogen group task, 1 NPC vs. any pathogen task, 1 VAP onset task, and 1 VAP cure task. Each task had 4 cross-validation folds, and we repeated model fine tuning 5 times for each fold with different random seed initializations to assess stability. We applied 3-layer MLP classification head to the CLS token, with output dimensions of 128, 64 and 1, using ReLU activation and layer normalization. During finetuning, we froze 3 out of 4 transformer layers and the token embeddings. We trained the models for 20 epochs using AdamW optimizer with constant learning rate.

After finetuning, we performed 16 forward passes on the test part of the cross-validation fold with random dropout enabled and averaged the resulting logits. We used these averaged logits to compute model performance metrics, including accuracy, precision, recall, binary F1 score, area under receiver operating characteristic (AUROC) curve, and area under precision-recall curve (AUPRC). Because the 5 models trained with different random initializations within each task and fold showed highly similar performance, their results were averaged for reporting.

To characterize feature importance and EHRformer behavior, we analyzed attention values across four cross-validation folds. For each correct prediction, we averaged the attention matrices over the attention heads. We then extracted the attention values corresponding to the CLS token in the final layer, and multiplied it by attention matrices of the preceding layers^68^. To obtain a feature-level attention score for each model, the attention value assigned to the CLS token was set to zero and the remaining attention values were renormalized across features to sum to one. Attention scores were averaged separately over positive and negative examples and then averaged across cross-validation folds.

### Comparative baseline using XGBoost

To establish a comparative baseline, we trained XGBoost^20^ models for each of the predictive tasks and each cross-validation fold. We used XGBoost with default parameters (objective=binary:logistic, eval_metric=logloss). The models were trained on the original numeric and categorical data for each patient-day, without tokenization described in the “Input representation” section for EHRformer). We interpreted feature importance in XGBoost models by computing Shapley values on correct predictions using the ‘interventional’ approach and summarizing them as the mean absolute value across all data splits^69,70^.

### BAL sample processing and flow cytometry

A subset of BAL samples was sent for flow cytometry (**Extended Data Fig. 1a**). BAL fluid samples were filtered through a 70-μm cell strainer, pelleted by centrifugation at 400 relative centrifugal force (rcf) for 10 min at 4 °C, followed by hypotonic lysis of red blood cells with 2 ml of PharmLyse (BD Biosciences) reagent for 2 min. Lysis was stopped by adding 13 ml of MACS buffer (Miltenyi Biotech). Cells were pelleted again and resuspended in 100 μl of a 1:10 dilution of Human TruStain FcX (BioLegend) in MACS buffer and a 10-μl aliquot was taken for counting using a K2 Cellometer (Nexcelom) with Acridine Orange (AO)/Propidium Iodide (PI) reagent. The cell suspension volume was adjusted so the concentration of cells was always <5 × 10^7^ cells ml^−1^, and the fluorophore-conjugated antibody cocktail was added in a 1:1 ratio. The following antibodies were used (antigen, clone, fluorochrome, manufacturer, catalog no., final dilution): CD4, RPA-T4, BUV395, BD, 564724, 1:40; CD19, HIB19, BUV395, BD, 740287, 1:40; CD25, 2A3, BUV737, BD, 564385, 1:20; CD56, NCAM16.2, BUV737, BD, 612766, 1:20; HLA-DR, L243, eFluor450, Thermo Fisher Scientific, 48-9952-42, 1:40; CD45, HI30, BV510, BioLegend, 304036, 1:20; CD15, HI98, BV786, BD, 563838, 1:20; CD3, SK7, PE, Thermo Fisher Scientific, 12-0036-42, 1:20; CD127, HIL-7R, PECF594, BD, 562397, 1:20; CD206, 19.2, PECy7, Thermo Fisher Scientific, 25-2069-42, 1:40; CD8, SK1, APC, BioLegend, 344721, 1:40; CD14, M5E2, APC, BioLegend, 301808, 1:40; and EpCAM, 9C4, APC, BioLegend, 324208, 1:40. After incubation at 4 °C for 30 min, cells were washed with 5 ml of MACS buffer, pelleted by centrifugation and resuspended in 500 μl of MACS buffer with 2 μl of SYTOX Green viability dye (Thermo Fisher Scientific). Cells were sorted on a FACS Aria III SORP instrument using a 100-μm nozzle. Cells were sorted into 300 μl of 2% bovine serum albumin (BSA) in Dulbecco’s phosphate-buffered saline and cryopreserved using the protocol by L. Mazutis^71^. Briefly, cells pelleted by centrifugation at 400 rcf for 5 min at 4 °C, resuspended in Bambanker freezing medium to ∼2,000 cells μl^−1^ concentration. Concentration was confirmed using a K2 Cellometer (Nexcelom) with AO/PI reagent using the ‘Immune cells low RBC’ program with default settings, and ∼40-μl aliquots were immediately frozen at −80 °C. Sample processing was performed in a BSL-2 facility using BSL-3 practices. Analysis of the flow cytometry data was performed using FlowJo v.10.7.1. using a sequential gating strategy reported in our previous publication^12,26^ (Extended Data Fig. 2a, b in Grant et al.) and reviewed by two investigators (S.S. and A.V.M.). Relative cell-type abundance was calculated as a percentage of all singlets/live/CD45+ cells.

### Single-cell RNA-seq sample sequencing

A subset of BAL samples was sent for single-cell RNA sequencing (**Extended Data Fig. 1a**). Samples for scRNA-seq were processed as previously described^12,26^. scRNA-seq was performed using Chromium Next GEM Single Cell 5′ reagents v.2 from 10x Genomics (protocol no. CG000331 Rev A). Immediately before loading 10x Genomics Chip K with Chromium Single Cell 5′-gel beads and reagents, aliquots of cryopreserved cells were retrieved from a −80 °C freezer, rapidly thawed in a water bath at 37 °C, gently mixed by pipetting and added to the mix at room temperature. The volume of the single-cell suspension was calculated using the above protocol from 10x Genomics, based on the concentration at the time of cryopreservation and aiming to capture 5,000–10,000 cells per library. Libraries were prepared according to the manufacturer’s protocol. After quality checks, scRNA-seq libraries were pooled and sequenced on a NovaSeq 6000 instrument.

### Single-cell RNA-seq data processing and cell type identification

We demultiplexed each sequencing run for single-cell RNA-seq libraries with the ‘cellranger mkfastq’ pipeline, generating separate FASTQ files for each single-cell RNA-seq library. We processed each library with the ‘cellranger count’ pipeline from cellranger v7.0.0 with ‘include_introns’ set to false and default options and aligned reads to a hybrid reference genome of human and SARS-CoV-2, as previously described^26^. We loaded all library count matrices with scanpy v1.9.5 and concatenated and filtered cells using the following criteria: genes detected in at most 50 out of 3,022,609 cells were removed, and cells with number of UMI below 200 or above 60,000 were removed. We selected 1000 highly-variable genes with the Pearson residuals method^72^ and additionally excluded cells with fewer than 10 UMIs of these genes. We did not include SARS-CoV-2 genes among the selected highly-variable genes. We integrated the data using the scVI model from scvi-tools v1.0.4 with the BAL sample identifier as the batch covariate. Next, we used scanpy leiden clustering over scVI latent space to cluster the cells, and removed clusters of low-quality cells and doublets. We subsequently repeated the workflow starting from highly-variable gene selection, followed by manual reclustering and cell type annotation based on canonical marker genes and published lung cell atlases^73^, identifying a total 28 cell types. Two patients withdrew from the study after sample processing, integration, and cell type identification.

We did not redo these analyses; however, we removed those samples from any subsequent analysis or presentation. Another four samples came from a separate Auxora study (IRB STU00214054), which did not include consent to share metadata included in this paper. Thus we did not perform any analyses that necessitated clinical data using these samples. The final object had 2,438,672 cells.

### Sample clustering based on cell type abundance

For each sample, we computed the percentage abundances of 7 immune cell types identified from flow cytometry (neutrophils, CD206^high^ macrophages, CD206^low^ macrophages, monocytes, CD8+ T cells, CD4+ T cells, T_reg_ cells). The obtained cell type abundance vectors were later transformed using element-wise square root transformation, z-scored, and clustered using the K-Means algorithm. To determine the number of clusters, we ran 30 different initializations of the K-Means algorithm for cluster numbers ranging from 5 to 30. Finally, we used the Fowkles-Mallows score-based consistency metric^74^ and selected 9 as the number of flow cytometry-based clusters.

We next performed sample clustering of cell type abundances based on the pre-processed and annotated single-cell data, focusing on 23 immune cell types out of the 28 annotated cell types. Because neutrophil populations are lost during the cryopreservation necessary to process and sequence most of these samples, we defined the abundance of neutrophils from the flow cytometry data and corrected the percentages of the other immune cell types from the scRNA-seq data as we previously described^12^. Using the same clustering and cluster number selection procedure as above, we chose 13 as the number of scRNA-seq-based clusters.

### Pseudobulk gene expression profiles

For several analyses of gene expression, including MOFA, PCA, differential gene expression analysis, and gene set variation analysis, we summarized gene expression for each sample and each annotated cell type from our scRNA-seq data. We required at least 50 cells to create a pseudobulk sample for a given sample and cell type. We summed integer counts for all cells across all genes, and saved this matrix for all cell types. We excluded several gene categories from the pseudobulk count matrices: mitochondrial genes, SARS-CoV-2 viral transcripts, ribosomal protein-coding genes (RPL and RPS families), and unannotated transcripts lacking canonical HGNC symbols (i.e., genes identified only by Ensembl or NCBI/RefSeq accession identifiers). These categories were excluded a priori because they reflect technical artifacts, viral rather than host signal, or transcripts without interpretable biological annotation, as we sought to elucidate established biological pathways as well as credential protein based biomarkers.

### Gene set variation analysis

We performed gene set variation analysis (GSVA)^19^ using 50 hallmark gene sets^75^ on the collection of our pseudobulk samples, including all cell types that had at least 50 samples. We used decoupler package^75^ to run GSVA with parameters kcdf=True and seed=1066.

### Unsupervised analysis using MOFA

To discover major axes of variation in gene expression among our large dataset of scRNA-seq samples, while accounting for differences in cell type composition between samples, we applied the Multi-Omic Factor Analysis (MOFA) framework^15^. With this approach, we modeled a set of latent factors for each sample that simultaneously explained variation in cell type proportions and average gene expression profiles of multiple cell types in that sample. We applied MOFA to 263 SCRIPT and 9 healthy volunteer samples.

### Pre-processing

To construct the cell type abundance sample data view for MOFA, we used the abundance of cell types for each sample from pre-processed and annotated single-cell data and percentage of neutrophils as detected by flow cytometry. We limited this analysis to 23 immune cell populations (B cells, CD4+ T cells, CD8 T+ cells, Classical monocytes-1 CCR2, Classical monocytes-2 IL1B, DC1, DC2, MRC1+C1QA+ AM, MRC1+C1QA–AM, Migratory DC, NUPR1+ AM, Non-classical monocytes, Interstitial macrophages, Plasma cells, Proliferating CD4+ T cells, Proliferating CD8+ T cells, Proliferating NUPR1+ AM, Proliferating γδT cells, Proliferating plasma cells, Tregs, γδT cells, pDC, Neutrophils), and adjusted fractions of all non-neutrophil cell types to sum to 1 – neutrophil fraction for each sample. Samples where neutrophil percentage from flow cytometry was not available were not included in this data view. Then we normalized cell type abundances to align distributions across data views for MOFA. Each sample was normalized to 10k, then log transformed with a pseudocount of 1 and quantile normalized the data to normal distribution.

Next, we selected and constructed gene expression data views for MOFA. First, we selected the cell types for which at least 50% (136) of samples had pseudobulks. This resulted in the list of 8 cell types: NUPR1+ AM, MRC1+C1QA+ AM, MRC1+C1QA– AM, Classical monocytes-2 IL1B, DC2, CD4+ T cells, CD8+ T cells, and γδT cells. Three samples had data in only 2 MOFA views (pseudobulk cell types or cell type abundance), and were excluded. For each cell type we used the pseudobulk gene expression matrix (as described above) and performed additional gene filtering: we removed genes encoding ribosomal proteins and genes not expressed robustly across samples, that is genes that were detected in less than 80% of samples. Next, we normalized pseudobulk gene expression matrices for each cell type to 10,000 total counts per each sample, followed by z-score transformation across genes and quantile normalization across samples.

Final dimensions of the MOFA input were 9 views × 269 samples × 80,788 features across all views. The samples were composed of 9 healthy, 25 NPC, 149 with any of our 9 pathogen groups for modeling, and 86 other samples that did not belong to any of these categories. See **Supplementary Table 38** for the breakdown of samples in the MOFA model across sample categories. The feature dimensions of each view were as follows:

Cell type abundance: 23 cell types

NUPR1+ AM: 11,467 genes

MRC1+ C1QA+ AM: 11,017 genes

MRC1+ C1QA– AM: 11,189 genes

Classical Monocytes-2 IL1B: 9,488 genes

DC2: 9,923 genes

CD4+ T cells: 9,521 genes

CD8+ T cells: 9,700 genes

γδT cells: 8,460 genes

### Running MOFA and model selection

We performed MOFA using the muon python package version 0.1.5, which runs the core mofapy2 MOFA code in the background (version 0.7.0). We used default parameters with the exception of center_groups=False as this was addressed by our preprocessing.

We chose a 10-factor model for the analyzed data, because increasing the number of factors only modestly improved variance explained by the model and did not identify additional clinically important associations. To improve factor interpretability for our dataset, we rotated the obtained MOFA factor matrix using Kabsch algorithm towards the following targets (factor maximization objectives)^76^:

1. Early SARS-CoV-2 samples
2. Samples with bacteria only detected
3. Samples with bacteria detected
4. Samples from immunocompromised patients
5. Samples from female patients
6. Samples from patients who died
7. Samples obtained when patient was on ECMO
8. Samples adjudicated to start CAP pneumonia episodes
9. VAP samples from VAP episodes that were cured by day 7 adjudication
10. Viral, bacterial or mixed samples

### Interpretation of MOFA factors

For each MOFA factor, there exists a weight for each of 80,765 genes across all cell type views. We applied GSEA^18^ to estimate enrichment of the 50 hallmark gene sets from MSigDB in gene lists ranked by corresponding weights for each factor and each cell type. We masked all genes coming from other cell types to ensure GSEA did not consider them a part of the gene set. We corrected estimated GSEA p-values for each MOFA factor for all gene sets and cell types with the FDR procedure. We assessed the significance of weights for cell type abundance features as p-values for z-scores of the weights in normal distributions fitted to each factor’s weights.

### Principal component analysis

Similar to MOFA, we ran PCA independently across annotated cell types. We used pseudobulk gene expression matrices for each cell type, obtained as described above. Next, we constructed a DESeq2 object with model expression ∼ sex and fitType set to ’local’ and applied DESeq2 variance-stabilizing transformation to obtain normalized pseudobulk counts.

For each cell type we selected the top 2,000 genes based on their variance across all normalized pseudobulk samples. Then we applied PCA to the resulting normalized count matrix, and analysed the top 10 PCs.

### Interpretation of MOFA factors and PCA principal components

To test associations between MOFA factors or PCA principal components with the sample covariates, we ran statistical tests separately for categorical and numerical covariates. For categorical covariates, we excluded missing values and performed a two-sided Mann-Whitney U test if there were just 2 categories or ANOVA if there were more. The tests were performed on factor values across groups defined by the covariate categories. For numerical covariates, we excluded missing values and ran Spearman correlation between factor values and covariate values. Next, we applied false discovery rate (FDR) correction for multiple testing (10 MOFA factors × number of categorical/numerical covariates), for categorical and numerical covariates separately. When multiple categorical variables were derived from the same underlying data (specifically, “discharge disposition” and “mortality”, as well as “pathogen groups” and “healthy/NPC/any pathogen”), only one representative variable was included in the FDR correction at a time and the procedure was repeated. In addition, for each categorical variable and each factor, we performed a pairwise two-sided Mann-Whitney U test to test for differences of factor values between each pair of values of the variable, followed by FDR correction. The same analysis was performed for PCA components.

### Differential gene expression analysis

To identify genes that are expressed differently between groups of samples, we used a modified pseudobulk method. This method demonstrates better consistency with the bulk approach in benchmarking studies^77^, compared to single cell based methods for testing of differentially expressed genes. Additionally, it avoids the issue of inflated p-values. We used pseudobulk gene count matrices as described above and constructed a DESeq2 object using all samples, split into 2 groups for comparison and the remaining samples for better estimation of gene dispersions. We used ‘∼ group + sex’ and fitType ‘local’ for the DESeq2 model. For each pairwise comparison, we then applied two independent filtering criteria: (i) we removed genes detected in less than 80% of pseudobulk samples in at least one of the two compared groups, where ‘detected’ was defined as a raw count ≥ 1, and (ii) DESeq2’s default independent filtering by baseMean. Genes failing these criteria were retained in the count matrix during model fitting so that they could contribute to size factor and gene dispersion estimation in DESeq2, and were excluded only at the FDR-adjustment and reporting stages. We used significance level of 0.05 for FDR.

We performed differential gene expression analysis between all pairs of these groups of samples: “healthy volunteers”, “NPC”, “early SARS-CoV-2”, “late SARS-CoV-2”, “bacterial” and “mixed infections”. These groups were defined from our infecting pathogen labels (see “Sample labels for machine learning” above). In addition, we compared gene expression between samples labeled as “gram+” and “gram–” (a union of “other gram–” and “*Pseudomonas*”); between samples labeled as “other gram–” and “*Pseudomonas*”; between samples from patients who were discharged vs. deceased, performing the comparisons for all samples, samples from patients within “early SARS-CoV-2”, “bacterial”, and “pathogen-negative” groups; between samples from the beginning of VAP episodes that lead to VAP cure within 7 days vs VAP failure to cure within 7 days; between samples from patients less than 65 years old vs. older; between samples from female vs. male patients; between samples from immunocompetent vs. immunocompromised patients; between samples from the bottom 50% vs. top when sorted by cumulative steroid dose administered to patients up to the day of sampling; and between samples from the first vs. last quartile when sorted by cumulative days of mechanical ventilation up to the day of sampling.

We also performed differential gene expression analysis to identify transcriptional changes associated with future VAP using serial BAL samples as follows. First, we identified patients with at least 2 BAL samples in our scRNA-seq dataset within 14 days of each other. For patients with 3 or more samples, we retained only the first pair of samples. We retained only those pairs of samples where the first sample belonged to an adjudicated CAP or HAP episode or was an NPC sample. We assigned pairs where the second sample belonged to a VAP episode to the “future VAP” group, and pairs where the second sample did not belong to a VAP episode to the “no future VAP” group. We kept other pairs of samples for gene dispersion estimation.

Next, we modeled gene expression in these pairs of samples with DESeq2 using the design ‘∼ group + group:timepoint + group:pair_id’ accounting for patient-specific baseline differences, group-level effects, and differences between the first and second sample within each pair. We compared gene expression in the first samples between groups, and between second and first samples within each group.

To summarize and interpret results of gene expression analysis, we applied GSEA^18^ to estimate enrichment of the 50 hallmark gene sets from MSigDB for each comparison ranked by DESeq2 reported log fold change for each cell type. We corrected estimated GSEA p-values for each comparison for all gene sets and cell types with the FDR procedure.

### Analysis using Geneformer

To assess transcriptional changes of lung immune cells predictive of pathogens or VAP onset or outcome at single-cell resolution, we used Geneformer^17^. We downloaded the V1-10M model (6 transformer layers, 2048 context size) model, which was originally published by Theodoris and colleagues, and finetuned this model for our defined classification tasks (see above). For visualization of Geneformer latent representations of our data we used the original Geneformer model without fine tuning.

To fine-tune, we fixed the bottom 4 layers and all token embeddings in Geneformer, and only allowed top 2 layers to change. Additionally, we increased model dropout in transformer layers to 0.2. For each classification task, we finetuned the model for 3 epochs on the labeled train data, propagating the labels from samples to all cells belonging to the samples. We downsampled the prevalent class to the minority class randomly. We used a cosine scheduler with 2e–5 learning rate, max_grad_norm=0.1 and 8 gradient accumulation steps. In total, we had 20 classification tasks for fine-tuning: 9 healthy volunteer samples vs. each pathogen group, 9 NPC samples vs. each pathogen group, early SARS-CoV-2 samples vs. bacterial samples, VAP cure in 7 days, and for each we finetuned 4 models for each cross-validation fold and repeated that 5 times for different random seed initializations, resulting in 400 models.

We evaluated our fine-tuned models by assessing their performance on the test part of the cross-validation fold. First, we ran forward passes of our models on test data 16 times with random dropout enabled and averaged the logits of the outputs, obtaining predictions for each cell. Then we averaged these predictions for each sample and each cell type or for each sample to obtain predictions on a sample level. Using sample-level predictions, we computed and reported AUROC and AUPRC for the test data.

Additionally, we used our fine-tuned models to classify samples labeled as “pathogen-negative”, “NPC”, “pathogens identified in a small number of samples” and samples from patients with post-acute sequelae of COVID-19, published previously^12^. To obtain predicted sample classification, we averaged predictions from all cells in a sample per model, and used average pathogen prediction probabilities across healthy volunteer samples vs. each pathogen group models with 0.5 threshold as pneumonia prediction, and early SARS-CoV-2 samples vs. bacterial only samples model with 0.5 threshold as viral vs. bacterial prediction.

## Data and code availability

Some data from patients in this cohort have been previously published^26,27,38,43^; clinical data for the cohort are available on Physionet.^22^ Code for EHR feature preprocessing is available at https://github.com/NUSCRIPT/carpediem_v1.9.

Code for analyses presented in this paper is available at https://github.com/NUPulmonary/2026_Pneumonia_endotypes. Interactive data exploration tools are available at https://sqlifts.fsm.northwestern.edu/public/pneumonia-endotypes. Flow cytometry data is available in **Supplementary Table 4**. Single-cell RNA-sequencing data will be made available upon publication.

## Acknowledgements

We thank all patients and volunteers who provided their samples and data for this study and their families. This research was supported in part through a generous gift from Kimberly Querrey and Louis A. Simpson. This research was supported by the Simpson Querrey Lung Institute for Translational Science (SQLIFTS), Northwestern University. This research was also supported by the computational resources and staff contributions provided for the Quest high-performance computing facility at Northwestern University, which is jointly supported by the Office of the Provost, the Office for Research and Northwestern University Information Technology. This research was also supported in part through the computational resources and staff contributions provided by the Genomics Compute Cluster, which is jointly supported by the Feinberg School of Medicine, the Center for Genetic Medicine and Feinberg’s Department of Biochemistry and Molecular Genetics, the Office of the Provost, the Office for Research and Northwestern Information Technology. The Genomics Compute Cluster is part of Quest, Northwestern University’s high-performance computing facility, with the purpose of advancing research in genomics. We thank Jackie Milhans, Alper Kinaci, Scott Coughlin, and all members of the Research Computing and Data Services team at Northwestern for their support. This work is dedicated to the benefit of all sentient beings. We are grateful to Stanisław Lem and his books for inspiration. Northwestern University Flow Cytometry Core Facility is supported by the National Cancer Institute Cancer Center support grant (no. P30 CA060553) awarded to the Robert H. Lurie Comprehensive Cancer Center. Cell sorting was performed on a BD FACS Aria SORP cell sorter purchased with the support of the National Institutes of Health (NIH, grant no. 1S10OD011996-01). Integrative genomic services were performed by the Metabolomics Core Facility at Robert H. Lurie Comprehensive Cancer Center of Northwestern University. Next-generation sequencing was performed with support from the Simpson Querrey Institute for Epigenetics.

N.S.M. was supported by the American Heart Association (grant no. 24PRE1196998) https://doi.org/10.58275/AHA.24PRE1196998.pc.gr.190609. M.M. was supported by “Excellence Initiative – Research University” Programme at the University of Warsaw, decision no. BOB-IDUB-622-645/2023. P.S. was supported by “Excellence Initiative – Research University” Programme at the University of Warsaw, decision no. BOB-IDUB-622-644/2023. A.I. was supported by “Excellence Initiative – Research University” Programme at the University of Warsaw, decision no. BOB-IDUB-622-643/2023. L.V.R. is supported by NIH/NIAID U19 AI135964. T.L.W. was supported by Gilead Sciences (award no. CO-US-540-6435) and the NIH (grant nos. U19AI135964, U19AI181102, and R21HD107571). R.G.W. is supported by NIH/NIAID U19 AI135964, RO1 AI158530; NHLBI RO1 HL149883, P01 HL154998; U01TR003528. G.R.S.B. is supported by the NIH (U19AI135964, P01AG049665, R01HL147575, P01HL071643, U54AG079754, R01HL145478, R01HL147290, R01HL173940, U19AI181102, and R01HL154686); the US Department of Veterans Affairs (I01CX001777); Simpson Querrey Lung Institute for Translational Sciences. C.A.G. is supported by NIH/NHLBI K23HL169815, a Parker B. Francis Opportunity Award, and an American Thoracic Society Unrestricted Grant. B.D.S. is supported by NIH awards R01HL149883, R01HL153122, P01HL154998, P01AG049665, U19AI135964, and U19AI181102. A.V.M. was supported by the NIH (grant nos. U19AI135964, P01AG049665, P01HL154998, P01HL169188, U19AI181102, R01HL153312, R01HL158139, and R01ES034350), and research grants from AbbVie and Merck.

## Author contributions

Conceptualization: N.S.M., T.W., R.G.W., G.R.S.B., L.M.-N., C.A.G., B.D.S., A.V.M. and E.S. Methodology: N.S.M., M.M., V.G., M.Ł., S.F., Al.P., B.P.-S., P.S., A.I., L.L., K.J.S., L.C., S.S., H.A.-V., R.G.W., L.M.-N., C.A.G., A.V.M. and E.S. Software: N.S.M., M.M., V.G., M.Ł., S.F., Al.P., B.P.-S., P.S., A.I., L.L., K.J.S., L.C. and An.P. Validation: N.S.M., M.M., V.G., S.F., Al.P., B.P.-S., P.S., A.I., L.L., K.J.S., L.C., L.V.R., R.G.W., L.M.-N., C.A.G., A.V.M. and E.S. Formal analysis: N.S.M., M.M., V.G., S.F., Al.P., B.P.-S., P.S., A.I., L.L., K.J.S., L.C., R.G.W., C.A.G., A.V.M. and E.S. Investigation: N.S.M., H.K.D., A.D., S.S., Z.L., H.A.-V., D.P., R.G.W., C.A.G., A.V.M. and E.S. Resources: N.S.M., H.K.D., A.D., Z.Y., S.S., Z.L., H.A.-V., D.P., C.O.P., N.R.N., R.M.T., R.G.W., G.R.S.B., L.M.-N., C.A.G., B.D.S., A.V.M. and E.S. Data curation: N.S.M., H.K.D., A.D., L.C., Z.Y., S.S., Z.L., H.A., An.P., C.O.P., N.R.N., R.M.T., R.G.W., G.R.S.B., L.M.-N., C.A.G., B.D.S., A.V.M. and E.S. Writing – all aspects: N.S.M., R.G.W., G.R.S.B., L.M.-N., C.A.G., B.D.S., A.V.M. and E.S. Writing – review & editing: C.O.P. Visualization: N.S.M., M.M., V.G., M.Ł., S.F., Al.P., B.P.-S., P.S., A.I., L.L., K.J.S. and L.C. Supervision: T.W., R.G.W., G.R.S.B., C.A.G., B.D.S., A.V.M. and E.S. Project administration: N.S.M., H.K.D., R.K.C., L.V.R., C.O.P., T.W., R.G.W., G.R.S.B., C.A.G., B.D.S., A.V.M. and E.S. Funding acquisition: L.V.R., C.O.P., T.W., R.G.W., G.R.S.B., B.D.S. and A.V.M.

## Extended data

**Extended Data Table 1.** Baseline demographic and clinical characteristics of healthy controls. . This table presents the demographic and clinical characteristics of the healthy control group. Continuous variables are summarized as median (interquartile range, IQR with Q1,Q3), and categorical variables are shown as number (percentage).

|  | Overall |
| --- | --- |
| n | 9 |
| Female, n (%) | 5 (55.6) |
| Male, n (%) | 4 (44.4) |
| Age, median [Q1,Q3] | 29.0 [24.0,33.0] |
| BMI, median [Q1,Q3] | 24.9 [23.4,31.7] |

**Extended Data Table 2.** Demographic and clinical outcomes for the patient cohort. . This table presents demographic characteristics and clinical outcomes for the full patient cohort. All variables were complete except for body mass index (BMI), which was missing for 6 patients. Discharge disposition was used to determine in-hospital mortality status; patients who died, were discharged to hospice or received a lung transplant during hospitalization were categorized as having had an unfavorable outcome. Racial groups with fewer than five individuals were classified as *‘Unknown or Not Reported’* to protect patient anonymity. Continuous variables are summarized as median (interquartile range), and categorical variables as number (percentage). Abbreviations: APS, Acute Physiology Score; BMI, Body Mass Index; CRRT, Continuous Renal Replacement Therapy; ECMO, Extracorporeal Membrane Oxygenation; ICU, Intensive Care Unit; LTACH, Long-Term Acute Care Hospital; NAT, Narrow Antibiotic Therapy; Q1, First Quartile; Q3, Third Quartile; scRNA-seq, Single-Cell RNA Sequencing; SOFA, Sequential Organ Failure Assessment; SNF, Skilled Nursing Facility; VAP, Ventilator-Associated Pneumonia.

|  | Overall | Non-pneumonia<br>Control | COVID-19 | Other pneumonia |
| --- | --- | --- | --- | --- |
| **n** | 690 | 99 | 167 | 424 |
| **Age (years), median [Q1,Q3]** | 62.0<br>[51.0,71.0] | 60.0 [50.0,70.0] | 61.0 [50.0,70.0] | 63.5 [52.0,72.0] |
| **Ethnicity, n (%)** |  |  |  |  |
| Hispanic or Latino | 141 (20.4) | 12 (12.1) | 64 (38.3) | 65 (15.3) |
| Not Hispanic or Latino | 517 (74.9) | 81 (81.8) | 98 (58.7) | 338 (79.7) |
| Unknown or Not Reported | 32 (4.6) | 6 (6.1) | 5 (3.0) | 21 (5.0) |
| **Female, n (%)** | 279 (40.4) | 49 (49.5) | 58 (34.7) | 172 (40.6) |
| **Race, n (%)** |  |  |  |  |
| Asian | 20 (2.9) | 4 (4.0) | 3 (1.8) | 13 (3.1) |
| Black or African American | 134 (19.4) | 18 (18.2) | 34 (20.4) | 82 (19.3) |
| Unknown or Not Reported | 127 (18.4) | 17 (17.2) | 48 (28.7) | 62 (14.6) |
| White | 409 (59.3) | 60 (60.6) | 82 (49.1) | 267 (63.0) |
| **Smoking Status, n (%)** |  |  |  |  |
| Former | 174 (25.2) | 27 (27.3) | 22 (13.2) | 125 (29.5) |
| Never | 282 (40.9) | 42 (42.4) | 69 (41.3) | 171 (40.3) |
| Unknown Smoking Status | 171 (24.8) | 22 (22.2) | 72 (43.1) | 77 (18.2) |
| Current | 63 (9.1) | 8 (8.1) | 4 (2.4) | 51 (12.0) |
| **Body Mass Index, median [Q1,Q3]** | 28.6<br>[24.2,33.8] | 27.5 [24.2,33.3] | 30.2 [26.7,35.8] | 27.6 [23.0,33.1] |
| **Immunocompromised, n (%)** | 208 (30.1) | 36 (36.4) | 31 (18.6) | 141 (33.3) |
| **Transferred from External Facility, n (%)** | 215 (31.2) | 32 (32.3) | 59 (35.3) | 124 (29.2) |
| **Admission APS Score, median [Q1,Q3]** | 91.0<br>[64.0,109.0] | 91.0 [63.0,106.0] | 93.0 [63.5,109.0] | 90.0 [65.0,109.0] |
| **Admission SOFA Score, median [Q1,Q3]** | 11.0 [8.0,14.0] | 11.0 [7.5,14.0] | 11.0 [8.0,13.0] | 11.0 [8.0,14.0] |
| **Developed VAP, n (%)** | 284 (41.2) | 16 (16.2) | 110 (65.9) | 158 (37.3) |
| **Cumulative NAT Score, median [Q1,Q3]** | 6.0 [-4.0,18.0] | 4.0 [-2.0,12.5] | -1.0 [-15.5,17.0] | 8.0 [0.0,20.0] |
| **Cumulative Steroid Dose, median [Q1,Q3]** | 770.0<br>[0.0,2067.5] | 650.0<br>[0.0,2815.0] | 1200.0<br>[75.0,2125.0] | 680.0 [0.0,2016.2] |
| **Received norepinephrine during admission, n (%)** | 597 (86.5) | 80 (80.8) | 156 (93.4) | 361 (85.1) |
| **Received hemodialysis during admission, n (%)** | 83 (12.0) | 11 (11.1) | 17 (10.2) | 55 (13.0) |
| **Received CRRT during admission, n (%)** | 201 (29.1) | 30 (30.3) | 43 (25.7) | 128 (30.2) |
| **Received ECMO during admission, n (%)** | 63 (9.1) | 5 (5.1) | 35 (21.0) | 23 (5.4) |
| **Has analyzed flow cytometry data, n (%)** | 482 (69.9) | 59 (59.6) | 144 (86.2) | 279 (65.8) |
| **Has analyzed scRNA-seq data, n (%)** | 170 (24.6) | 24 (24.2) | 81 (48.5) | 65 (15.3) |
| **Total ICU Days, median [Q1,Q3]** | 16.0 [8.0,28.8] | 10.0 [5.5,18.5] | 26.0 [15.5,38.5] | 13.0 [7.0,23.0] |
| **Number of ICU Admissions, median [Q1,Q3]** | 1.0 [1.0,1.0] | 1.0 [1.0,1.0] | 1.0 [1.0,1.0] | 1.0 [1.0,2.0] |
| **Underwent Tracheostomy, n (%)** | 179 (25.9) | 14 (14.1) | 73 (43.7) | 92 (21.7) |
| **Total Intubation Days, median [Q1,Q3]** | 11.0 [5.2,24.0] | 7.0 [4.0,13.0] | 24.0 [12.0,36.5] | 10.0 [5.0,19.0] |
| **Discharge Disposition, n (%)** |  |  |  |  |
| **Home** | 147 (21.3) | 27 (27.3) | 39 (23.4) | 81 (19.1) |
| **Rehab** | 113 (16.4) | 11 (11.1) | 21 (12.6) | 81 (19.1) |
| **LTACH** | 78 (11.3) | 5 (5.1) | 22 (13.2) | 51 (12.0) |
| **SNF** | 41 (5.9) | 6 (6.1) | 8 (4.8) | 27 (6.4) |
| **Hospice** | 24 (3.5) | 8 (8.1) |  | 16 (3.8) |
| **Death** | 287 (41.6) | 42 (42.4) | 77 (46.1) | 168 (39.6) |
| **Mortality, n (%)** | 311 (45.1) | 50 (50.5) | 77 (46.1) | 184 (43.4) |

**Extended Data Table 3.** Demographic and clinical outcomes for the patient cohort who had flow cytometry data analyzed. . This table presents demographic characteristics and clinical outcomes for the subset of patients with at least one BAL sample that was analyzed using flow cytometry. All variables were complete except for body mass index (BMI), which was missing for 6 patients. Discharge disposition was used to determine in-hospital mortality status; patients who died, were discharged to hospice or received a lung transplant during hospitalization were categorized as having had an unfavorable outcome. Racial groups with fewer than five individuals were classified as *‘Unknown or Not Reported’* to protect patient anonymity. Continuous variables are summarized as median (interquartile range), and categorical variables as number (percentage). Abbreviations: APS, Acute Physiology Score; BAL, bronchoalveolar lavage; BMI, Body Mass Index; CRRT, Continuous Renal Replacement Therapy; ECMO, Extracorporeal Membrane Oxygenation; ICU, Intensive Care Unit; LTACH, Long-Term Acute Care Hospital; NAT, Narrow Antibiotic Therapy; Q1, First Quartile; Q3, Third Quartile; scRNA-seq, Single-Cell RNA Sequencing; SOFA, Sequential Organ Failure Assessment; SNF, Skilled Nursing Facility; VAP, Ventilator-Associated Pneumonia.

|  | Overall | Non-pneumonia<br>Control | COVID-19 | Other<br>pneumonia |
| --- | --- | --- | --- | --- |
| **n** | 482 | 59 | 144 | 279 |
| **Age (years), median [Q1,Q3]** | 62.0 [50.0,71.0] | 63.0 [47.0,70.5] | 60.5 [50.0,69.0] | 63.0 [52.0,71.0] |
| **Ethnicity, n (%)** |  |  |  |  |
| Hispanic or Latino | 110 (22.8) | 6 (10.2) | 59 (41.0) | 45 (16.1) |
| Not Hispanic or Latino | 352 (73.0) | 49 (83.1) | 80 (55.6) | 223 (79.9) |
| Unknown or Not Reported | 20 (4.1) | 4 (6.8) | 5 (3.5) | 11 (3.9) |
| **Female, n (%)** | 192 (39.8) | 28 (47.5) | 49 (34.0) | 115 (41.2) |
| **Race, n (%)** |  |  |  |  |
| White | 276 (57.3) | 34 (57.6) | 67 (46.5) | 175 (62.7) |
| Black or African American | 98 (20.3) | 13 (22.0) | 29 (20.1) | 56 (20.1) |
| Asian | 12 (2.5) | 1 (1.7) | 2 (1.4) | 9 (3.2) |
| Unknown or Not Reported | 96 (19.9) | 11 (18.6) | 46 (31.9) | 39 (14.0) |
| **Smoking Status, n (%)** |  |  |  |  |
| Never Smoked | 188 (39.0) | 22 (37.3) | 56 (38.9) | 110 (39.4) |
| Former Smoker | 120 (24.9) | 16 (27.1) | 21 (14.6) | 83 (29.7) |
| Current Smoker | 40 (8.3) | 5 (8.5) | 2 (1.4) | 33 (11.8) |
| Unknown | 134 (27.8) | 16 (27.1) | 65 (45.1) | 53 (19.0) |
| **Body Mass Index (kg/m<sup>2</sup>), median [Q1,Q3]** | 28.9 [24.6,33.8] | 27.0 [24.3,32.2] | 30.3 [26.9,35.8] | 27.8 [23.2,33.2] |
| **Immunocompromised, n (%)** | 131 (27.2) | 19 (32.2) | 23 (16.0) | 89 (31.9) |
| **Transferred from External Facility, n (%)** | 162 (33.6) | 20 (33.9) | 55 (38.2) | 87 (31.2) |
| **Admission APS Score, median [Q1,Q3]** | 91.0 [64.0,109.0] | 93.0 [64.0,110.0] | 94.5 [64.8,110.0] | 89.0 [64.0,108.0] |
| **Admission SOFA Score, median [Q1,Q3]** | 11.0 [8.0,14.0] | 11.0 [7.0,14.0] | 11.0 [9.0,13.0] | 11.0 [8.0,14.0] |
| **Developed VAP, n (%)** | 208 (43.2) | 9 (15.3) | 100 (69.4) | 99 (35.5) |
| **Cumulative NAT Score, median [Q1,Q3]** | 6.5 [-4.0,19.0] | 3.0 [-1.5,10.5] | -0.5 [-16.0,17.0] | 9.0 [1.0,22.0] |
| **Cumulative Steroid Dose, median [Q1,Q3]** | 750.2 [0.0,2065.0] | 625.0 [0.0,2522.5] | 1175.0 [0.0,2222.5] | 660.0 [0.0,1810.0] |
| **Received norepinephrine during admission, n (%)** | 423 (87.8) | 44 (74.6) | 137 (95.1) | 242 (86.7) |
| **Received hemodialysis during admission, n (%)** | 54 (11.2) | 5 (8.5) | 14 (9.7) | 35 (12.5) |
| **Received CRRT during admission, n (%)** | 144 (29.9) | 14 (23.7) | 38 (26.4) | 92 (33.0) |
| **Received ECMO during admission, n (%)** | 59 (12.2) | 4 (6.8) | 35 (24.3) | 20 (7.2) |
| **Has analyzed scRNA-seq data, n (%)** | 158 (32.8) | 18 (30.5) | 79 (54.9) | 61 (21.9) |
| **Total ICU Days, median [Q1,Q3]** | 17.0 [8.0,30.0] | 8.0 [5.0,19.0] | 27.5 [16.0,39.0] | 14.0 [8.0,25.0] |
| **Number of ICU Admissions, median [Q1,Q3]** | 1.0 [1.0,1.0] | 1.0 [1.0,1.0] | 1.0 [1.0,1.0] | 1.0 [1.0,2.0] |
| **Underwent Tracheostomy, n (%)** | 142 (29.5) | 9 (15.3) | 68 (47.2) | 65 (23.3) |
| **Total Intubation Days, median [Q1,Q3]** | 12.0 [6.0,27.0] | 6.0 [3.0,11.5] | 26.5 [13.0,37.2] | 10.0 [5.0,20.0] |
| **Discharge Disposition, n (%)** |  |  |  |  |
| Home | 103 (21.4) | 18 (30.5) | 35 (24.3) | 50 (17.9) |
| Rehab | 77 (16.0) | 9 (15.3) | 16 (11.1) | 52 (18.6) |
| LTACH | 54 (11.2) | 4 (6.8) | 18 (12.5) | 32 (11.5) |
| SNF | 23 (4.8) | 2 (3.4) | 4 (2.8) | 17 (6.1) |
| Hospice | 15 (3.1) | 4 (6.8) |  | 11 (3.9) |
| Death | 210 (43.6) | 22 (37.3) | 71 (49.3) | 117 (41.9) |
| Mortality, n (%) | 225 (46.7) | 26 (44.1) | 71 (49.3) | 128 (45.9) |

**Extended Data Table 4.** Demographic and clinical outcomes for the patient cohort who had scRNA-seq data analyzed. . This table presents demographic characteristics and clinical outcomes for the subset of patients with at least one BAL sample that was analyzed using scRNA-seq. All variables were complete except for body mass index (BMI), which was missing for 6 patients. Discharge disposition was used to determine in-hospital mortality status; patients who died, were discharged to hospice or received a lung transplant during hospitalization were categorized as having had an unfavorable outcome. Racial groups with fewer than five individuals were classified as *‘Unknown or Not Reported’* to protect patient anonymity. Continuous variables are summarized as median (interquartile range), and categorical variables as number (percentage). Abbreviations: APS, Acute Physiology Score; BAL, bronchoalveolar lavage; BMI, Body Mass Index; CRRT, Continuous Renal Replacement Therapy; ECMO, Extracorporeal Membrane Oxygenation; ICU, Intensive Care Unit; LTACH, Long-Term Acute Care Hospital; NAT, Narrow Antibiotic Therapy; Q1, First Quartile; Q3, Third Quartile; scRNA-seq, Single-Cell RNA Sequencing; SOFA, Sequential Organ Failure Assessment; SNF, Skilled Nursing Facility; VAP, Ventilator-Associated Pneumonia.

|  | Overall | Non-pneumonia<br>Control | COVID-19 | Other pneumonia |
| --- | --- | --- | --- | --- |
| **n** | 170 | 24 | 81 | 65 |
| **Age (years), median [Q1,Q3]** | 59.0 [49.2,68.8] | 56.5 [47.0,67.0] | 59.0 [50.0,70.0] | 59.0 [47.0,67.0] |
| **Ethnicity, n (%)** |  |  |  |  |
| Hispanic or Latino | 45 (26.5) | 2 (8.3) | 29 (35.8) | 14 (21.5) |
| Not Hispanic or Latino | 116 (68.2) | 20 (83.3) | 49 (60.5) | 47 (72.3) |
| Unknown or Not Reported | 9 (5.3) | 2 (8.3) | 3 (3.7) | 4 (6.2) |
| **Female, n(%)** | 70 (41.2) | 14 (58.3) | 29 (35.8) | 27 (41.5) |
| **Male** | 100 (58.8) | 10 (41.7) | 52 (64.2) | 38 (58.5) |
| **Race, n (%)** |  |  |  |  |
| White | 93 (54.7) | 12 (50.0) | 43 (53.1) | 38 (58.5) |
| Black or African American | 31 (18.2) | 5 (20.8) | 15 (18.5) | 11 (16.9) |
| Asian | 3 (1.8) |  | 1 (1.2) | 2 (3.1) |
| Unknown or Not Reported | 43 (25.3) | 7 (29.2) | 22 (27.2) | 14 (21.5) |
| **Smoking Status, n (%)** |  |  |  |  |
| Never Smoked | 69 (40.6) | 6 (25.0) | 41 (50.6) | 22 (33.8) |
| Former Smoker | 36 (21.2) | 6 (25.0) | 10 (12.3) | 20 (30.8) |
| Current Smoker | 9 (5.3) | 3 (12.5) |  | 6 (9.2) |
| Unknown | 56 (32.9) | 9 (37.5) | 30 (37.0) | 17 (26.2) |
| **Body Mass Index (kg/m<sup>2</sup>), median [Q1,Q3]** | 29.0 [25.5,34.8] | 29.2 [23.8,36.4] | 29.1 [26.5,34.3] | 28.6 [24.8,34.7] |
| **Immunocompromised, n (%)** | 47 (27.6) | 9 (37.5) | 19 (23.5) | 19 (29.2) |
| **Transferred from External Facility, n (%)** | 55 (32.4) | 7 (29.2) | 29 (35.8) | 19 (29.2) |
| **Admission APS Score, median [Q1,Q3]** | 90.0 [60.2,107.2] | 92.0 [54.8,106.2] | 94.0 [64.0,109.0] | 86.0 [61.0,103.0] |
| **Admission SOFA Score, median [Q1,Q3]** | 11.0 [8.0,13.0] | 8.5 [6.8,13.2] | 11.0 [8.0,13.0] | 10.0 [8.0,13.0] |
| **Developed VAP, n (%)** | 92 (54.1) | 4 (16.7) | 58 (71.6) | 30 (46.2) |
| **Cumulative NAT Score, median [Q1,Q3]** | 3.0 [-6.0,19.0] | 2.5 [-4.5,11.5] | 1.0 [-15.0,16.0] | 7.0 [-3.0,23.0] |
| **Cumulative Steroid Dose, median [Q1,Q3]** | 1280.0 [200.0,2903.8] | 1075.0 [312.5,3435.0] | 1350.0 [150.0,2450.0] | 1250.0 [200.0,3950.0] |
| **Received norepinephrine during admission, n (%)** | 157 (92.4) | 19 (79.2) | 78 (96.3) | 60 (92.3) |
| **Received hemodialysis during admission, n (%)** | 18 (10.6) | 3 (12.5) | 9 (11.1) | 6 (9.2) |
| **Received CRRT during admission, n (%)** | 51 (30.0) | 7 (29.2) | 20 (24.7) | 24 (36.9) |
| **Received ECMO during admission, n (%)** | 29 (17.1) | 2 (8.3) | 20 (24.7) | 7 (10.8) |
| **Has analyzed flow cytometry data, n (%)** | 158 (92.9) | 18 (75.0) | 79 (97.5) | 61 (93.8) |
| **Has analyzed scRNA-seq data, n (%)** | 170 (100.0) | 24 (100.0) | 81 (100.0) | 65 (100.0) |
| **Total ICU Days, median [Q1,Q3]** | 23.0 [13.0,37.8] | 10.0 [4.8,18.5] | 29.0 [19.0,40.0] | 21.0 [11.0,38.0] |
| **Number of ICU Admissions, median [Q1,Q3]** | 1.0 [1.0,1.0] | 1.0 [1.0,1.0] | 1.0 [1.0,1.0] | 1.0 [1.0,1.0] |
| **Underwent Tracheostomy, n (%)** | 68 (40.0) | 4 (16.7) | 42 (51.9) | 22 (33.8) |
| **Total Intubation Days, median [Q1,Q3]** | 20.5 [9.0,33.0] | 5.5 [2.0,13.0] | 27.0 [13.0,37.0] | 16.0 [8.0,32.0] |
| **Discharge Disposition, n (%)** |  |  |  |  |
| **Home** | 33 (19.4) | 8 (33.3) | 16 (19.8) | 9 (13.8) |
| **Rehab** | 22 (12.9) | 3 (12.5) | 9 (11.1) | 10 (15.4) |
| **LTACH** | 18 (10.6) | 1 (4.2) | 10 (12.3) | 7 (10.8) |
| **SNF** | 5 (2.9) |  | 1 (1.2) | 4 (6.2) |
| **Hospice** | 3 (1.8) | 1 (4.2) |  | 2 (3.1) |
| **Death** | 89 (52.4) | 11 (45.8) | 45 (55.6) | 33 (50.8) |
| **Mortality, n (%)** | 92 (54.1) | 12 (50.0) | 45 (55.6) | 35 (53.8) |

**Extended Data Table 5.**
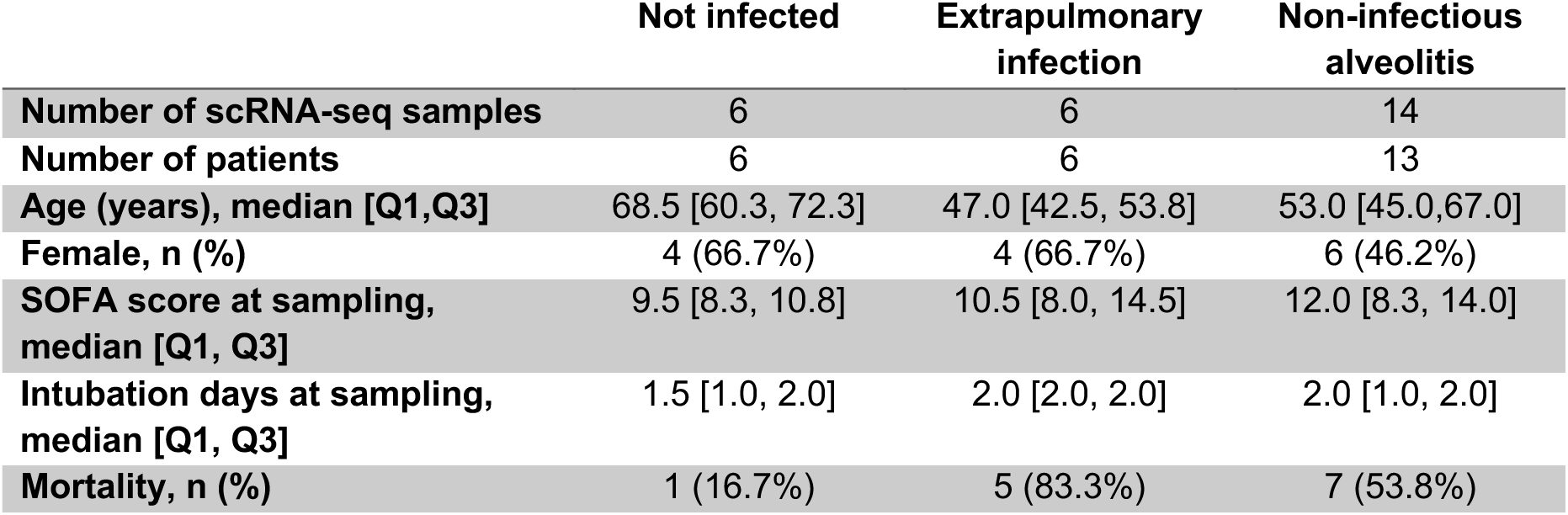
Demographic and clinical variables for adjudicated subtypes of NPC samples. Demographic, outcome and hospitalization information for the subset of samples and patients who had samples adjudicated as non-pneumonia controls (NPC) and profiled with single-cell RNA-sequencing (scRNA-seq). Abbreviations: Q1, First Quartile; Q3, Third Quartile; scRNA-seq, Single-Cell RNA Sequencing; SOFA, Sequential Organ Failure Assessment.

|  | **Not infected** | **Extrapulmonary infection** | **Non-infectious alveolitis** |
| --- | --- | --- | --- |
| **Number of scRNA-seq samples** | 6 | 6 | 14 |
| **Number of patients** | 6 | 6 | 13 |
| **Age (years), median [Q1,Q3]** | 68.5 [60.3, 72.3] | 47.0 [42.5, 53.8] | 53.0 [45.0,67.0] |
| **Female, n (%)** | 4 (66.7%) | 4 (66.7%) | 6 (46.2%) |
| **SOFA score at sampling, median [Q1, Q3]** | 9.5 [8.3, 10.8] | 10.5 [8.0, 14.5] | 12.0 [8.3, 14.0] |
| **Intubation days at sampling, median [Q1, Q3]** | 1.5 [1.0, 2.0] | 2.0 [2.0, 2.0] | 2.0 [1.0, 2.0] |
| **Mortality, n (%)** | 1 (16.7%) | 5 (83.3%) | 7 (53.8%) |

**Extended Data Fig. 1:**
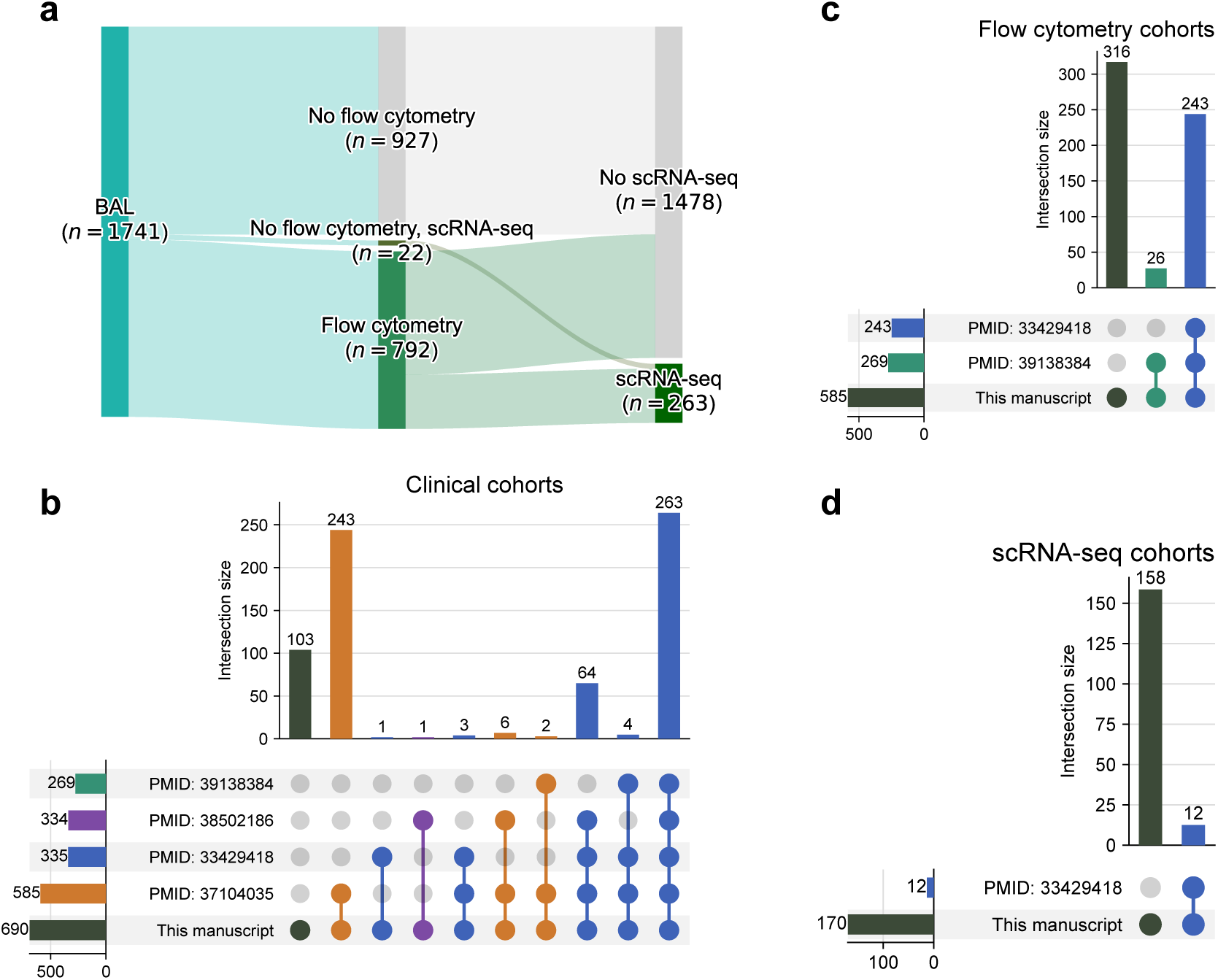
a. Alluvial plot describing the analysis of bronchoalveolar lavage (BAL) samples from the cohort with flow cytometry and single cell RNA-sequencing (scRNA-seq). **b.** Upset plots indicating overlap of clinical data from patients in the SCRIPT cohort included in this analysis with those included in previous analyses published from this cohort. **c.** Upset plots indicating overlap of flow cytometry data from patients in the SCRIPT cohort included in this analysis with those included in previous analyses published from this cohort. **d.** Upset plots indicating overlap of scRNA-seq data from patients in the SCRIPT cohort included in this analysis with those included in previous analyses published from this cohort.

**Extended Data Fig. 2:**
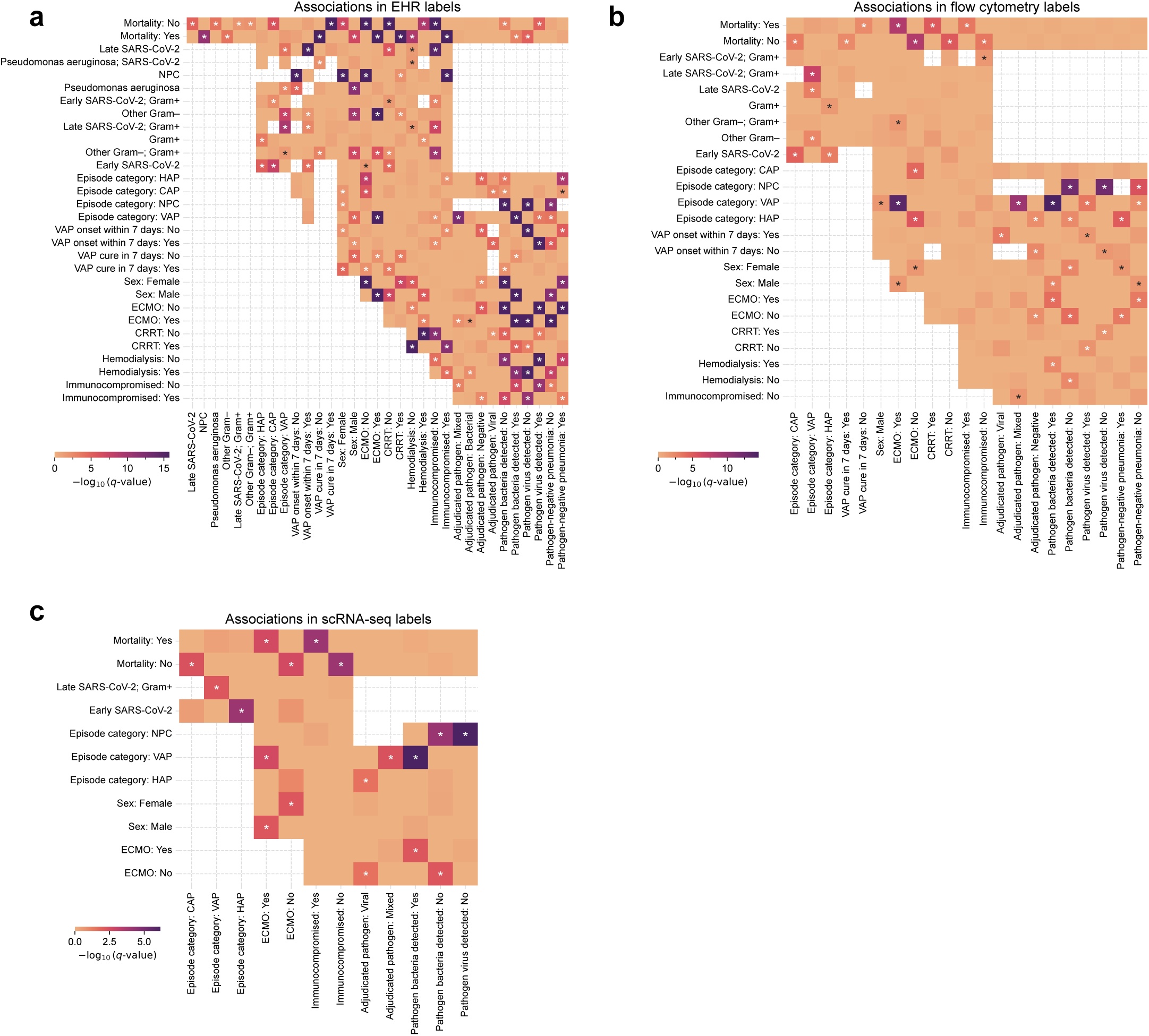
a. Heatmap of significant pre-existing associations in the EHR cohort, as tested with right-tailed hypergeometric tests with FDR correction. **b.** Heatmap of significant pre-existing associations in the flow cytometry cohort, as tested with right-tailed hypergeometric tests with FDR correction. **c.** Heatmap of significant pre-existing associations in the scRNA-seq cytometry cohort, as tested with right-tailed hypergeometric tests with FDR correction.

**Extended Data Fig. 3:**
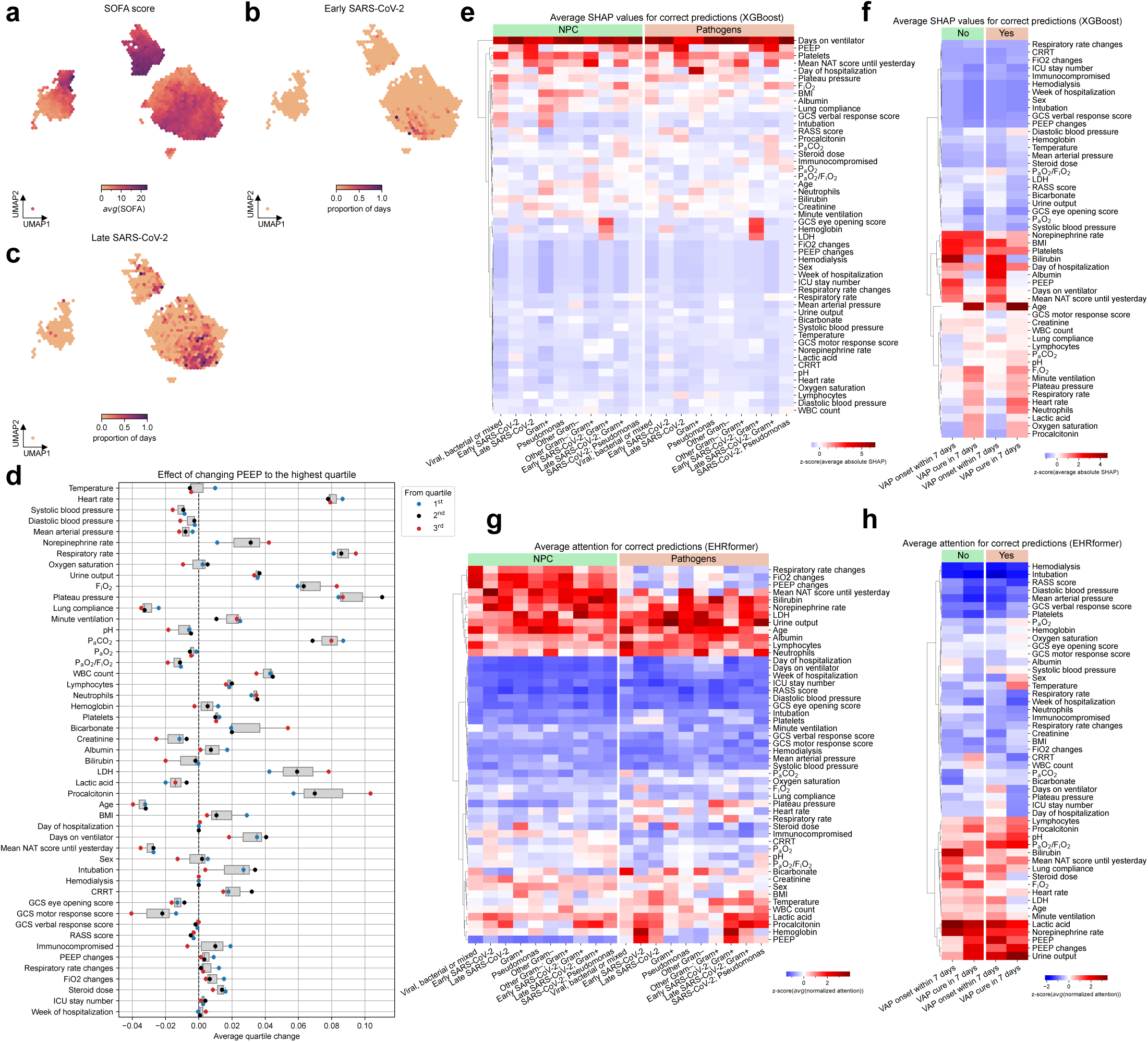
a. Hexbin plot of UMAP of EHRformer embeddings of ICU patient-day clinical features, colored by the sequential organ failure assessment (SOFA) score (a numeric measure of illness severity in patients with critical illness between 0–24) in each hexagon. **b.** colored by the proportion of patient-days coming from early SARS-CoV-2 episodes (≤ 48 hours after intubation) in each hexagon. **c.** colored by the proportion of patient-days from late SARS-CoV-2 episodes (> 48 hours after intubation). **d.** Effect (average quartile change; x-axis) of simulated perturbation in which the value of PEEP was set to the highest quartile, and quartile predictions were obtained from EHRformer for each other clinical feature (y-axis). **e.** Average SHapley Additive exPlanations (SHAP) values for pathogen predictions from the application of XGBoost to clinical features. **f.** Average SHAP values for predicting the onset of or cure after ventilator associated pneumonia (VAP) from the application of XGBoost to clinical features. **g.** Average attention for correct pathogen predictions from the application of EHRFormer to clinical features. **h.** Average attention for correct prediction of the onset of or cure after VAP from the application of EHRFormer to clinical features.

**Extended Data Fig. 4:**
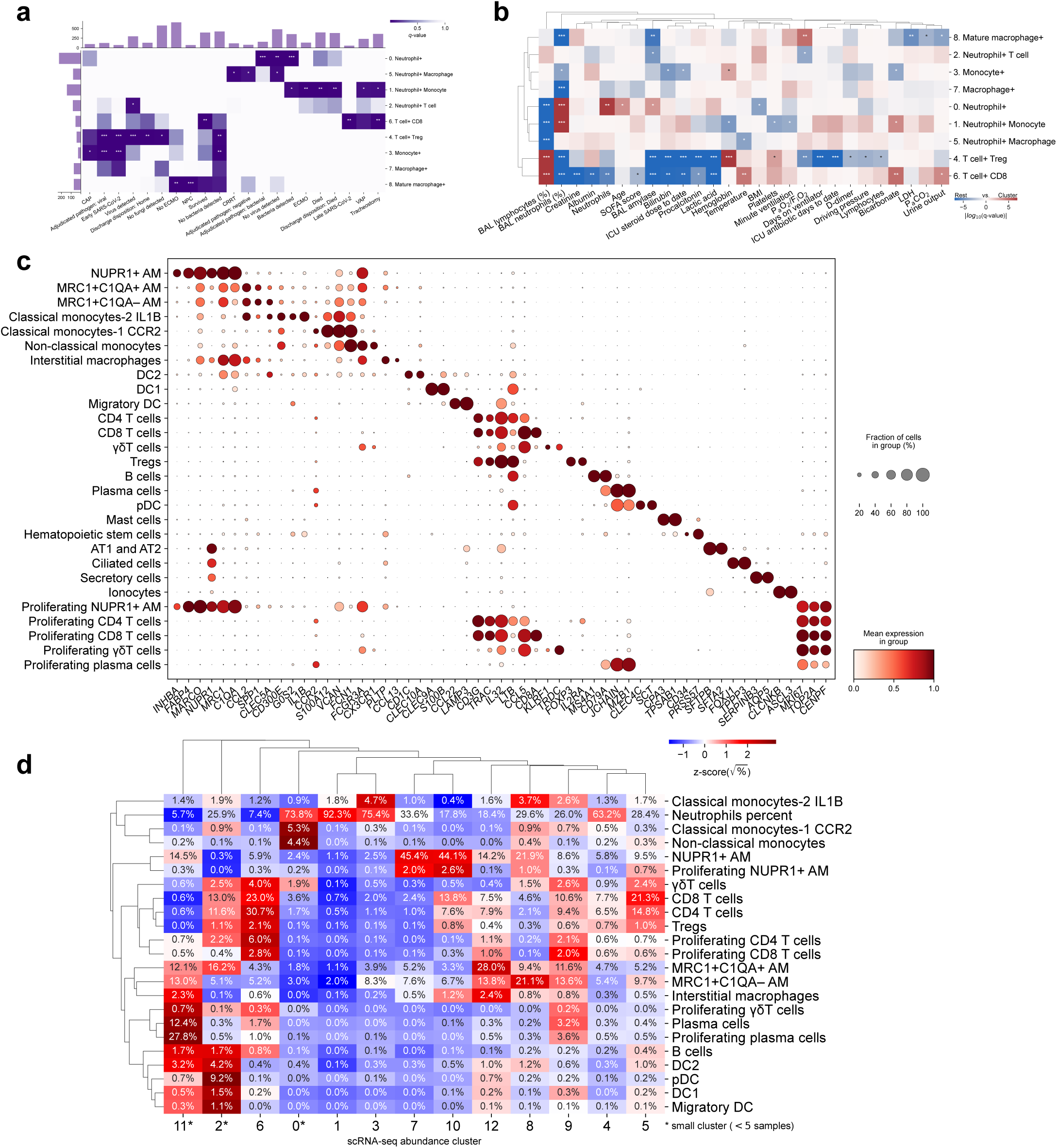
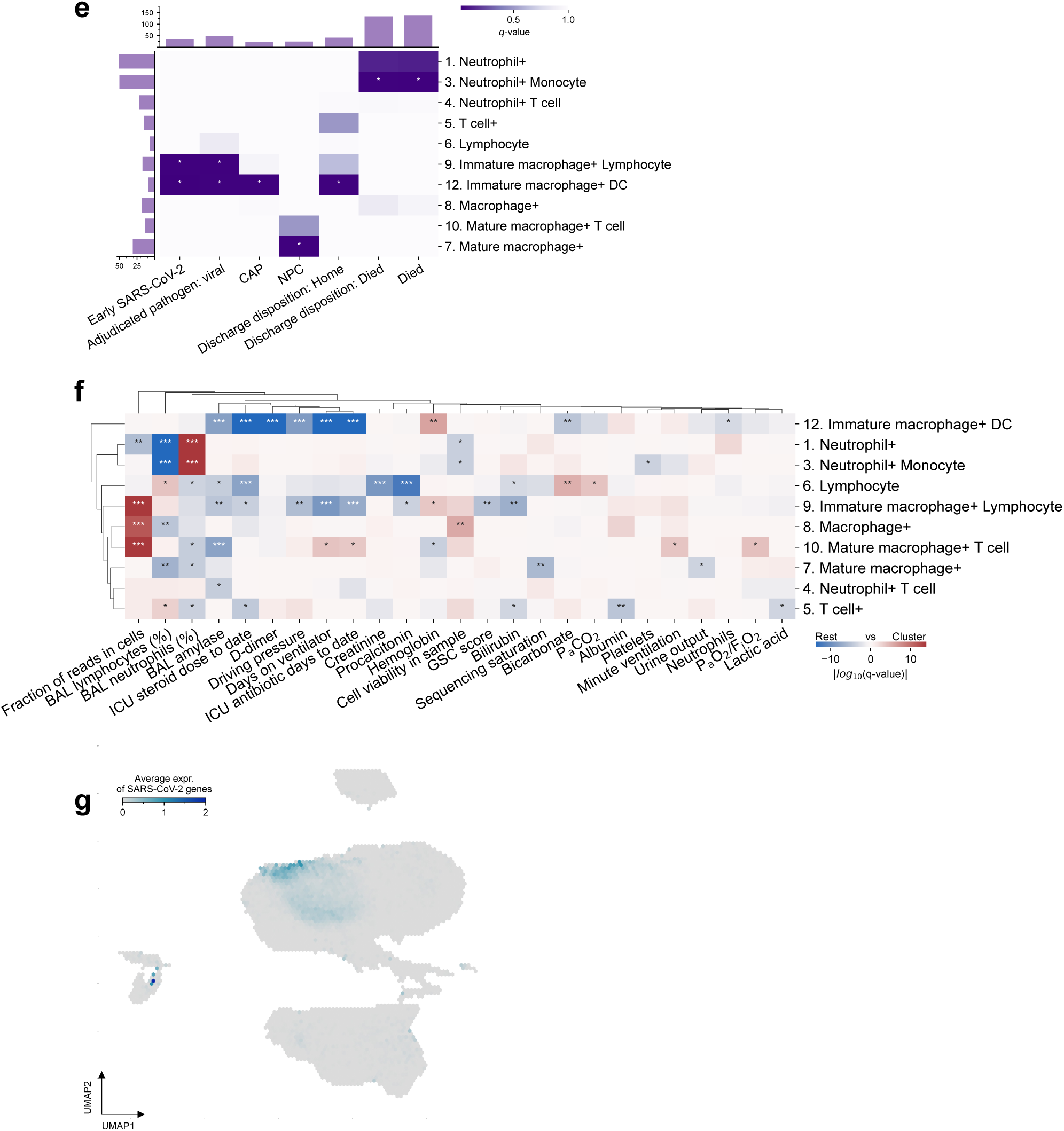
a. Associations between cell abundance clusters (rows) defined from flow cytometry and categorical pathogen groups and clinical features (columns). Pathogen groups and clinical features with significant associations are shown (right-tailed hypergeometric tests with FDR correction; * *q*-value < 0.05, ** *q*-value < 0.01, *** *q*-value < 0.001). “Adjudicated pathogen”: expert adjudication of pathogen cause of pneumonia episode (viral, bacterial, mixed, negative [pneumonia in the absence of identified pathogen]). “Bacteria detected”, “Virus detected” or “Fungi detected”: a positive PCR, culture or fungal culture of BAL for a bacterium, virus or fungi, respectively. “VAP”: ventilator-associated pneumonia. “Early SARS-CoV-2” or “Late SARS-CoV-2”: BAL sample positive for SARS-CoV-2 collected ≤ or > 48 hours after intubation, respectively. “NPC”: non-pneumonia control. “VAP cure in 7 days”: clinical adjudication of resolved VAP episodes resolved within 7 days. “ECMO”: the requirement for extracorporeal membrane oxygenation. “CRRT”: the requirement for continuous renal replacement therapy. **b.** Associations between cell abundance clusters (rows) defined from flow cytometry and continuous clinical features (columns). Clinical features with significant associations are shown (Welch’s unequal variances *t*-test of values inside vs. outside cluster with FDR correction; * *q*-value < 0.05, ** *q*-value < 0.01, *** *q*-value < 0.001). Color indicates if values inside the cluster are higher (red) or lower (blue) than outside. “BAL”: bronchoalveolar lavage, “SOFA”: sequential organ failure assessment score, “BMI”: body mass index, “PaO2/FiO2”: the partial pressure of oxygen in the arterial blood /fractional inspired concentration of oxygen, “Driving pressure” = plateau pressure – positive end expiratory pressure (PEEP), “LDH”: serum lactate dehydrogenase, “PaCO2”: the partial pressure of CO2 in the arterial blood. **c.** Representative marker genes used to annotate cell clusters identified by scRNA-seq. **d.** Normalized average percentage of the cell populations (rows) in each cluster as detected using scRNA-seq (columns). Clusters were named based on one or two most abundant cell populations. **e.** Associations between cell abundance clusters (rows) defined from scRNA-seq and categorical pathogen groups and clinical features (columns) which were significant for flow cytometry clusters. Pathogen groups and clinical features with significant associations are shown (right-tailed hypergeometric test; * *q*-value < 0.05, ** *q*-value < 0.01, *** *q*-value < 0.001). **f.** Associations between cell abundance clusters (rows) defined from scRNA-seq data and continuous clinical features (columns). Clinical features with significant associations are shown (Welch’s unequal variances *t*-test of values inside vs. outside cluster with FDR correction; * *q*-value < 0.05, ** *q*-value < 0.01, *** *q*-value < 0.001). “GCS”: Glasgow Coma Scale. **g.** Hexbin plot of UMAP of scRNA-seq data colored by average normalized expression of all SARS-CoV-2 transcripts.

**Extended Data Fig. 5:**
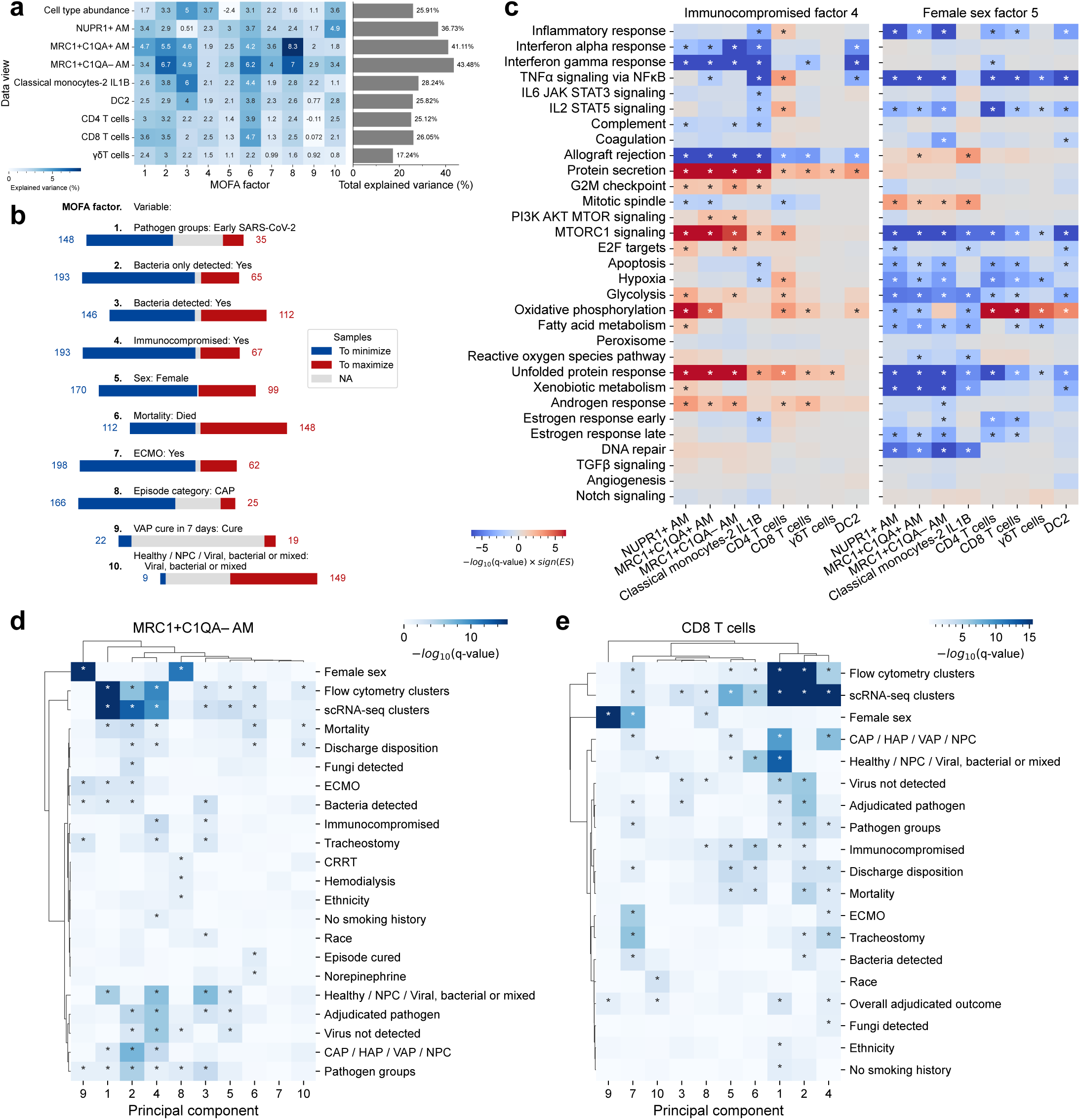
a. Explained variance attributed to each data view (rows) for each factor (columns) as a percentage of the total variance captured by the MOFA analysis. The bar graph on the right indicates the total variance explained for each data view by all 10 factors. **b.** Segmented bar chart showing the number of samples that were used to maximize (red) or minimize (blue) MOFA rotations. Blank bar indicates samples that do not meet either criteria. **c.** GSEA enrichment of hallmark gene sets (rows) in gene lists ranked by MOFA weights per each cell-type view (columns) for factors “immunocompromised factor 4” and “female sex factor 5”. * indicates significant enrichment using GSEA after FDR correction. **d.** Associations between discrete clinical features and the top 10 principal components of MRC1+C1QA– AM cell pseudobulk expression. Pseudobulk profiles were computed by averaging per-gene expression across all MRC1+C1QA– AM cells within each sample, and PCA was applied to the resulting samples-by-genes matrix. * indicates significant association as determined by two-sided Mann-Whitney U test (for categories with 2 levels) or ANOVA test with FDR correction. **e.** Associations between the first 10 principal components of pseudobulk CD8 T cell gene expression (samples × genes matrix) and discrete clinical features. * indicates significant association as determined by two-sided Mann-Whitney U test (for categories with 2 levels) or ANOVA test with FDR correction.

**Extended Data Fig. 6:**
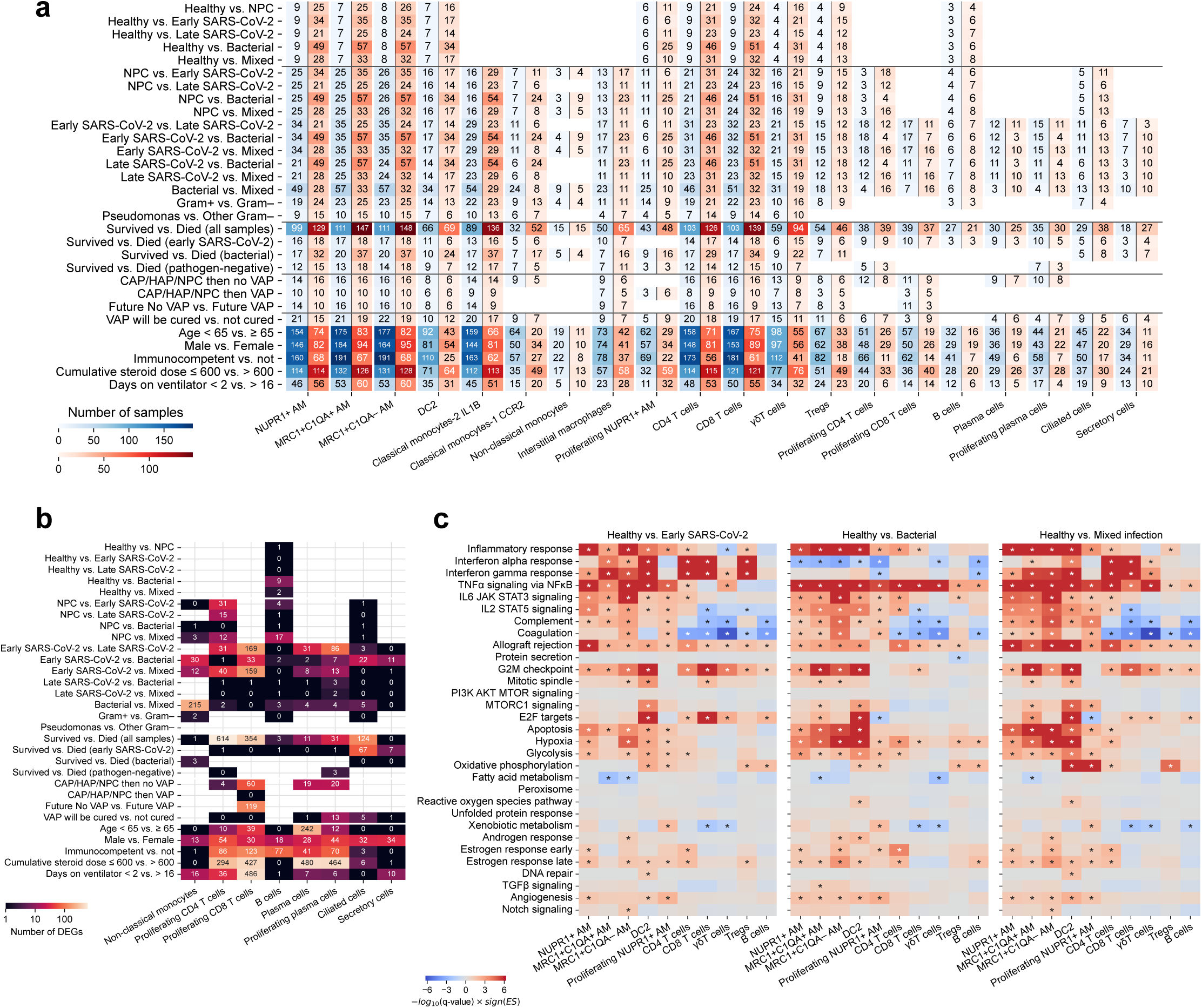
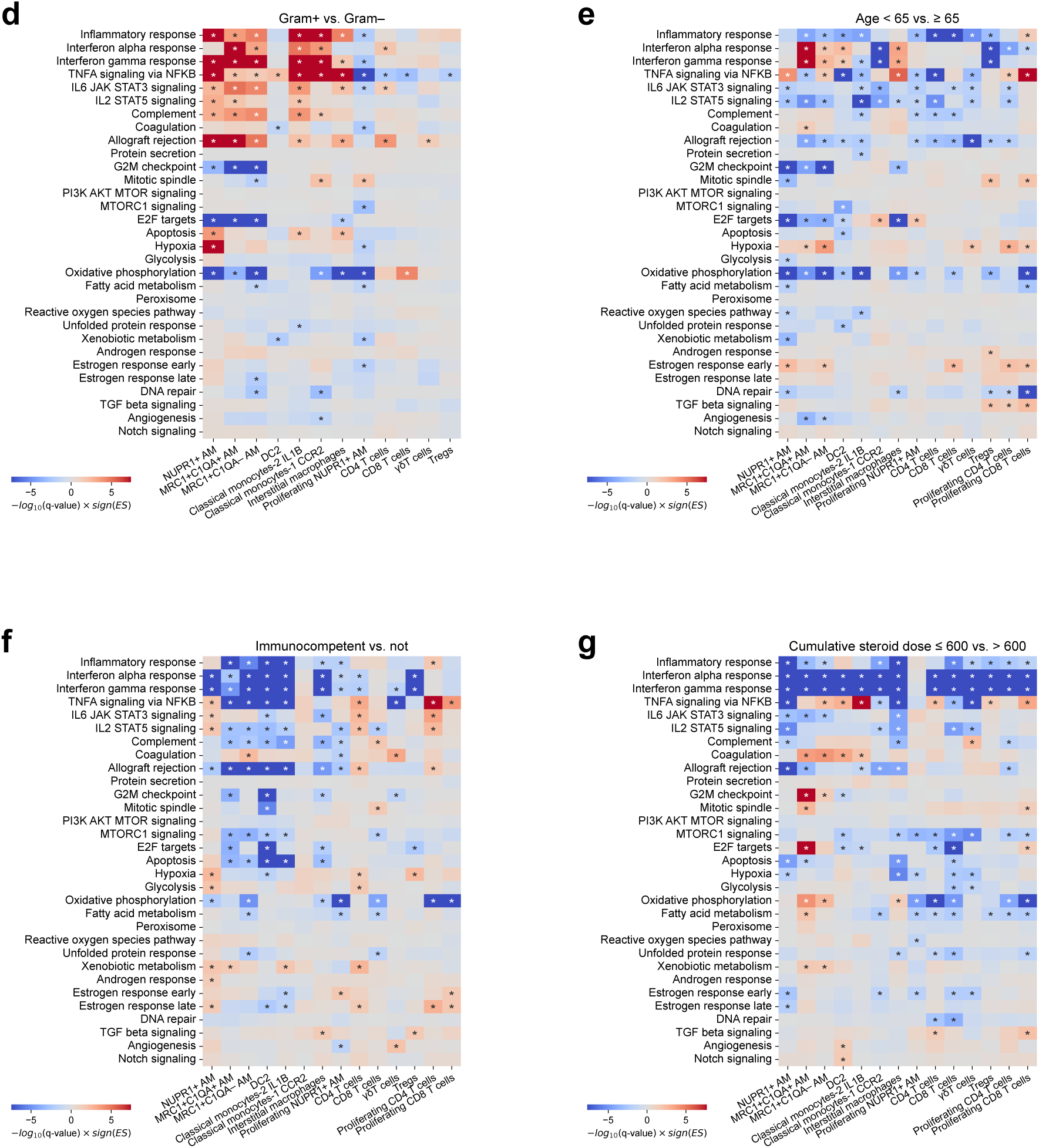
a. Number of samples for each comparison and cell type. **b.** Number of DEGs for each comparison for cell types not reported on Fig. 4a **c.** GSEA enrichment of hallmark gene sets (rows) in gene lists ranked by log-fold change per each cell-type (columns) for comparisons of sample groups “Healthy” vs. “early SARS-CoV-2” (left), “bacterial” (middle) and “mixed infection” (right) samples. * indicates significant enrichment using GSEA after FDR correction. **d.** GSEA enrichment of hallmark pathways (rows) for each cell type (columns) for comparisons of sample groups: “gram+” vs. “gram–” infected samples **e.** GSEA enrichment of hallmark gene sets (rows) for each cell type (columns) for comparisons of sample groups: patients younger than 65 years vs. 65 and older **f.** GSEA enrichment of hallmark gene sets (rows) for each cell type (columns) for comparisons of sample groups: immunocompetent vs. not immunocompetent patients **g.** GSEA enrichment of hallmark gene sets (rows) for each cell type (columns) for comparisons of sample groups: samples from patients who received ≤ 600 mg of hydrocortisone equivalent vs. > 600 mg during their ICU stay at the time of sampling

**Extended Data Fig. 7:**
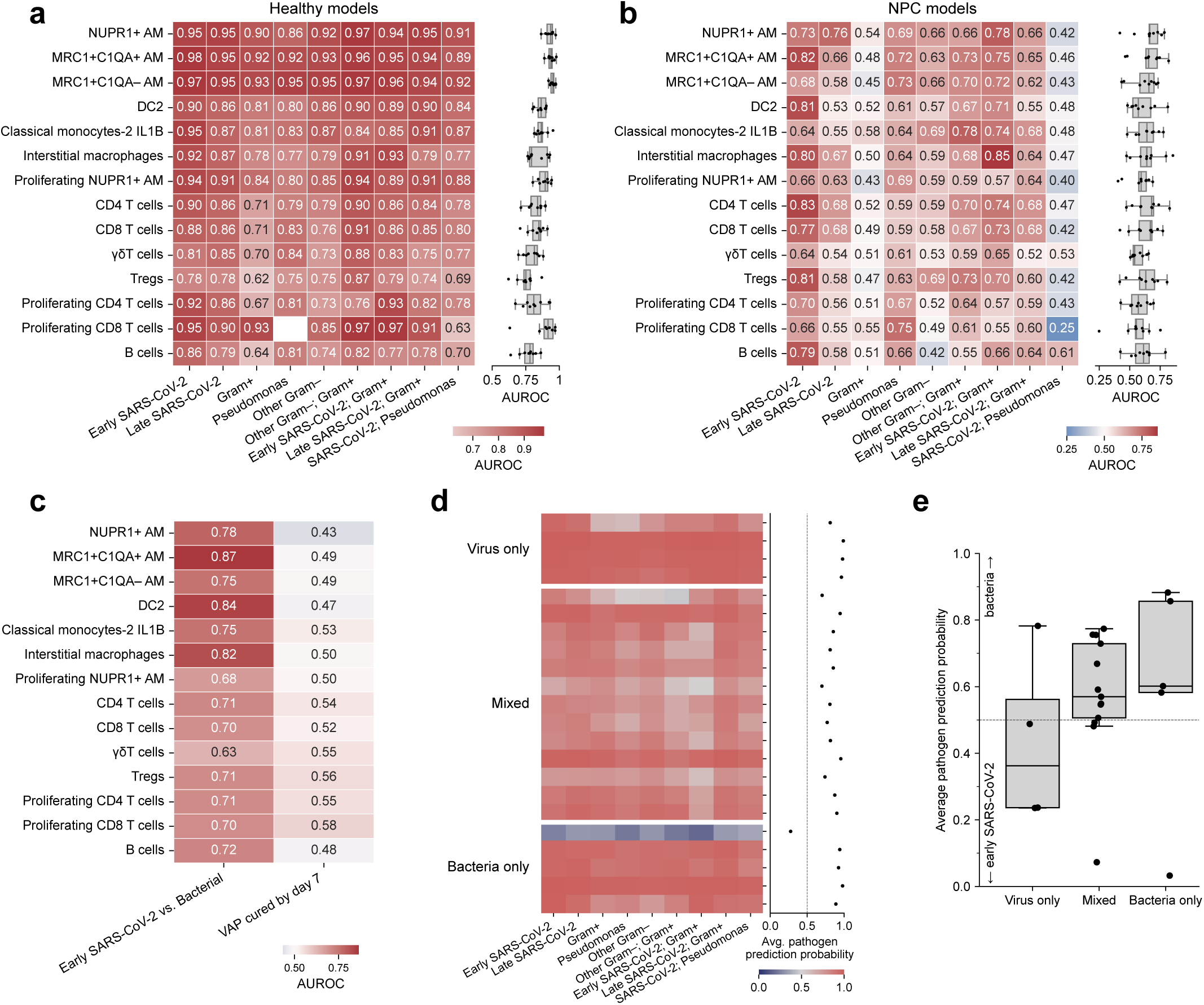
a. Heatmap of per-cell-type (rows) AUROC scores from fine-tuned Geneformer models classifying healthy versus pathogen-exposed samples (columns). **b.** Heatmap of per-cell-type (rows) AUROC scores from fine-tuned Geneformer models classifying NPC vs. pathogen groups models (columns). **c.** Heatmap of per-cell-type (rows) AUROC scores from fine-tuned Geneformer models classifying “early SARS-CoV-2” vs. “bacterial” and “VAP cure in 7 days” models (columns). **d.** Left: heatmap of average pathogen prediction probability from fine-tuned Geneformer models classifying healthy vs. pathogen groups models, for 22 samples (rows) in which pneumonia was attributed to a pathogen that was underrepresented in our dataset and were excluded from the primary analysis. Samples are grouped by pathogens detected in the sample: viral, bacterial or mixed viral and bacterial. Right: box-and-whisker plots of the mean predicted probability of detecting any pathogen across groups. **e.** Box-and-whiskers plot showing probability of classification as early SARS-CoV-2 pneumonia or bacterial pneumonia for the samples in (d).

## Supplementary information

Supplementary Videos are also available at https://sqlifts.fsm.northwestern.edu/public/pneumonia-endotypes/

**Supplementary Video 1. Demonstration of interactive visualization tool to explore MOFA results. Text for Supplementary Video 1.**

This movie explains how to use online tools to explore data from multi-factor omics analysis or MOFA applied to our single cell RNA sequencing data. The page can be accessed through the link to the webtool provided in the manuscript. For this example, we will demonstrate exploration of MOFA factor 5, which is associated with female sex. We chose this example as the mechanisms underlying the reduced risk of poor outcomes from pneumonia in women are incompletely understood.

When accessing MOFA, one sees two interactive heatmaps. The heatmap on the left identifies associations between MOFA factors and selected categorical metadata included in the dataset. The heatmap on the right identifies associations between MOFA factors and selected continuous variables in the metadata. Stars in a cell indicate a significant association as determined by hypergeometric test or Spearman correlation for categorical and numerical data, respectively, followed by FDR correction for multiple testing.

Focusing first on the heatmap on the left, one sees the 10 factors from MOFA on the bottom of the heatmap. For this example we are interested in Factor 5, which is significantly associated with female sex and a reduced requirement for tracheostomy, a measure of illness severity. When one clicks on the intersection of Factor 5 with sex (only significant associations can be explored), a boxplot will appear below with a per sample score for factor 5 segregated by sex. Hovering over the bar indicating significance will provide you with the numerical FDR corrected q value. A similar procedure shows the reduced association with tracheostomy.

One can now return to the heatmap and click on the text for Factor 5. This will open a new heatmap which provides the GSEA values associated with the gene rankings in factor 5 according to cell type as defined using single cell RNA sequencing. Blue on this heatmap means negatively enriched in female sex (or enriched in male sex). These data can be explored further by clicking on the significant association. For example, genes involved in MTORC signaling are negatively enriched for female sex across all cell populations. One can examine the leading edge genes in this pathway for each cell type by clicking on the corresponding cell. For example, the leading edge genes in the MTORC1_SIGNALING PATHWAY in immature monocyte derived alveolar macrophages (annotated as MRC1+C1QA- macrophages) include several genes implicated in the integrated stress response, for example BCAT-1, PPP1R15A or GADD34 and DDIT3 or CHOP. The leading edge genes in any other pathway or cell type can be explored the same way.

We will now go back to the main page for MOFA and turn our attention to the heatmap on the right. Here Factor 5 is associated with a higher driving pressure, possibly reflecting a higher plateau pressure, associated with improved respiratory system compliance or reduced requirements for PEEP in women, which can be visualized as a correlation plot by clicking on the cell. Clicking on the Factors on the heatmap on the right takes you to the same GSEA plots we discussed earlier.

**Supplementary Video 2. Demonstration of interactive visualization tool to explore gene expression analysis results.**

**Text for Supplementary Video 2**.

This video demonstrates the use of a webtool to access pseudobulk gene expression data generated from the single cell RNA sequencing object. The webtool can be accessed through the link provided in the manuscript.

The landing page for differential gene expression analysis includes a heatmap with comparisons between selected groupings of samples represented as rows and cell types from our single cell object as columns labeled on the bottom. Blank cells are those for which less than 3 samples contain a minimum of 50 cells from that cell type. Each cell includes a number representing the number of differentially expressed genes within that cell population for the indicated comparison. For example, in MRC1+C1QA+ alveolar macrophages there are 2473 differentially expressed genes between samples adjudicated as NPC compared to those with early SARS CoV-2 pneumonia, defined as a sample collected within 48 hours of intubation.

Hovering the mouse over an individual cell will display metadata on the right hand part of the screen. This metadata includes the number of samples in each group, the number of samples in the background (used to estimate gene variance, but not part of the comparison) and the total number of pseudobulk samples in the dataset for that cell type. The number of differentially expressed genes is indicated, which matches the number in the cell, subsetted into the number of genes that have increased and decreased expression. For example, the number 2473 represents the genes that are differentially expressed in MRC1+C1QA+ alveolar macrophages between NPC vs. Early SARS CoV-2 pneumonia and is based on a comparison of 25 samples adjudicated as NPC and 35 samples adjudicated as early SARS CoV-2 pneumonia. This number includes 1356 genes that are increased in NPC samples and 1117 genes that are increased in early SARS CoV-2 samples.

On the left side of the heatmap, each row label is a link to another heatmap. We will click on npc versus early sars cov 2 pneumonia. On this page, GSEA values are displayed for the indicated comparison by cell type, with red and blue labels indicating the direction of enrichment as indicated in the banner. For example there is enrichment of interferon-alpha and -gamma responses across multiple cell populations in early SARS-CoV-2 samples relative to NPC. By going back to the main page, we can contrast this with npc versus bacteria where we see enrichment for genes involved in TNFα signaling via NF-κB across multiple cell populations Staying on this GSEA heatmap and hovering over an individual cell displays the leading-edge genes from the indicated pathway identified as increased or decreased for that cell and comparison. For example, hovering over TNFα signaling via NF-κB in MRC1+C1QA+ alveolar macrophages reveals upregulation of recognizable NF-κB target genes like TNF, IL6 and REL. We can then click on this to lock that list on the top part of the screen and open a new box below it. The new box compares the differentially expressed genes in MRC1+C1QA+ alveolar macrophages with any other cell you choose to however over. For example, there are a large number of shared genes in the TNFα via NF-κB pathway between MRC1+C1QA+ and MRC1+C1QA- alveolar macrophages but fewer in CD8 T cells.

Going back to the main landing page, each cell in the heatmap links to a new page. The example of NPC vs Early SARS CoV-2 pneumonia in MRC1+C1QA+ alveolar macrophages will again be used to illustrate the use of this feature. Clicking on this cell opens a volcano plot that shows the genes with increased expression in samples from patients with early SARS CoV-2 pneumonia in red and increased expression in samples from episodes adjudicated as non-pneumonia controls in blue. On the right,the hallmark gene sets that are significantly different in this comparison are listed. Clicking on any dot shows the corresponding gene information on the right of the screen. The search bar shown below can be used to search for an individual gene. For example, CCL8 and RIGi (encoded by DDX58) were identified as the two most highly upregulated proteins in serum from people with influenza A infection compared to other infectious diseases in the Human Protein Atlas. Using the search function we see both are significantly upregulated in MRC1+C1QA+ alveolar macrophages from patients with early SARS CoV-2 pneumonia relative to non-pneumonia controls. Finally, clicking on the Hallmark Gene Sets on the right side of the screen displays the leading edge genes that are both expressed in MRC1+C1QA+ alveolar macrophages and are significantly upregulated or downregulated in the comparison. For example, 64 leading edge genes in the interferon-alpha response pathway are upregulated and 3 are downregulated in samples from patients with early SARS-CoV-2 pneumonia compared to those from non-pneumonia controls. These genes are also highlighted in the volcano plot on the left side of the screen.

**Supplementary Video 3. Demonstration of interactive visualization tool to explore gene set variation analysis results.**

**Text for Supplementary Video 3**.

This movie explains how to use online tools to explore data for Gene Set Variation Analysis (GSVA) applied to our single cell RNA sequencing data. The page can be accessed through the link to the webtool provided in the manuscript. This tool provides an interactive format to explore sample level enrichment in GSVA hallmark pathways as a function of selected metadata within our dataset.

This tool is useful for identifying similarities or differences in gene expression as a function of the infecting pathogen or clinical features and outcomes and includes samples from pneumonia pathogens that were excluded from our primary analyses because they were underrepresented in our dataset.

The landing for the GSVA analysis includes all of the 267 samples in the single cell RNA sequencing data as an individual column for the cell types shown in the box on the right side of the screen with the default opening to mature resident alveolar macrophages annotated as NUPR1+ alveolar macrophages in our dataset.

Clicking on a cell type takes you to a different heat map for that cell type. Please note that the number of samples represented as columns will vary by cell type as samples with fewer than 50 cells of that type were excluded from the analysis. Please also note that pathogen labels are blank for BAL samples collected outside of adjudicated episodes. . For this example, we will focus on more mature monocyte derived alveolar macrophages, annotated as MRC1+C1QA+ alveolar macrophages in our dataset.

There are two sets of rows. Those on the bottom, which are labeled on the right side of the figure represent the 50 hallmark processes in the GSVA analysis. Those on the top, which are labeled on the left side of the figure represent the selected metadata.Sample group, mortality, pathogen groups, flow-cytometry clusters,scRNA-seq clusters, and sex are selected by default.

If you hover to the left of each metadata label a small waffle will appear. If you click on that waffle it will arrange all of the columns according to the categories within that metadata field. For example clicking the waffle next to mortality will separate samples from patients who lived or died, clicking the waffle next to sample group will provide labels for all of the columns. Clicking on the tools option allows you to perform a variety of clustering functions. For example, we can perform k-means clustering with 5 clusters. This results in a new row of metadata. Clicking on the adjacent waffle will separate the columns by the k means cluster.

To see additional metadata, click on the View tab and then click on options. You will see three tabs on top, click on Annotations. Clicking on id in the Row annotations will provide the Hallmark Gene Set ID visible on the right of the heatmap. Clicking on column annotations will allow you to select additional metadata. For this example, we will choose all. These then appear to the left of the heatmap.

To give an example of how this tool can be used to explore sample level heterogeneity, we will focus on non-pneumonia controls. For this example, we will reopen the GSVA explorer from the landing page to reset to the default settings. We begin by clicking the View, options tab, which can also be accessed using the shortcut on the right of the screen. We select all metadata annotations for this example. We now filter the data to focus only on NPC samples by clicking the filter button and then selecting NPC, including all three subheadings. Next we widen the columns for readability using the options tab or selecting fit columns to window. We now click on the waffle next to NPC category, which sorts the columns by extrapulmonary infection, non-infectious alveolitis, and not infected. We see generally lower levels of inflammatory pathways like TNFα signaling via NF-κB in NUPR1+AM in the not infected group.

## List of Supplementary Tables

**Supplementary Table 1**. Numbers of samples for each value of each label for EHR, flow cytometry and scRNA-seq analysis groups.

**Supplementary Table 2**. Performance of XGBoost in predicting pathogen classification, VAP onset and cure, measured by area under receiver operating characteristic curve (AUROC) and area under precision-recall curve (AUPRC).

**Supplementary Table 3**. Performance of EHRformer in predicting pathogen classification, VAP onset and cure, measured by area under receiver operating characteristic curve (AUROC) and area under precision-recall curve (AUPRC).

**Supplementary Table 4**. Flow-cytometry measurements of proportion of 7 lung immune cell types in BAL samples.

**Supplementary Table 5**. Results of gene expression analysis for “Healthy” vs. “NPC” comparison for all tested cell types.

**Supplementary Table 6**. Results of gene expression analysis for “Healthy” vs. “Early SARS-CoV-2” comparison for all tested cell types.

**Supplementary Table 7**. Results of gene expression analysis for “Healthy” vs. “Late SARS-CoV-2” comparison for all tested cell types.

**Supplementary Table 8**. Results of gene expression analysis for “Healthy” vs. “Bacterial” comparison for all tested cell types.

**Supplementary Table 9**. Results of gene expression analysis for “Healthy” vs. “Mixed” comparison for all tested cell types.

**Supplementary Table 10**. Results of gene expression analysis for “NPC” vs. “Early SARS-CoV-2” comparison for all tested cell types.

**Supplementary Table 11**. Results of gene expression analysis for “NPC” vs. “Late SARS-CoV-2” comparison for all tested cell types.

**Supplementary Table 12**. Results of gene expression analysis for “NPC” vs. “Bacterial” comparison for all tested cell types.

**Supplementary Table 13**. Results of gene expression analysis for “NPC” vs. “Mixed” comparison for all tested cell types.

**Supplementary Table 14**. Results of gene expression analysis for “Early SARS-CoV-2” vs. “Late SARS-CoV-2” comparison for all tested cell types.

**Supplementary Table 15**. Results of gene expression analysis for “Early SARS-CoV-2” vs. “Bacterial” comparison for all tested cell types.

**Supplementary Table 16**. Results of gene expression analysis for “Early SARS-CoV-2” vs. “Mixed” comparison for all tested cell types.

**Supplementary Table 17**. Results of gene expression analysis for “Late SARS-CoV-2” vs. “Bacterial” comparison for all tested cell types.

**Supplementary Table 18**. Results of gene expression analysis for “Late SARS-CoV-2” vs. “Mixed” comparison for all tested cell types.

**Supplementary Table 19**. Results of gene expression analysis for “Bacterial” vs. “Mixed” comparison for all tested cell types.

**Supplementary Table 20**. Results of gene expression analysis for “Gram+” vs. “Gram–” comparison for all tested cell types.

**Supplementary Table 21**. Results of gene expression analysis for “Pseudomonas” vs. “Other Gram–” comparison for all tested cell types.

**Supplementary Table 22**. Results of gene expression analysis for “Survived” vs. “Died” comparison among all samples for all tested cell types.

**Supplementary Table 23**. Results of gene expression analysis for “Survived” vs. “Died” comparison among “early SARS-CoV-2” samples for all tested cell types.

**Supplementary Table 24**. Results of gene expression analysis for “Survived” vs. “Died” comparison among “bacterial” samples for all tested cell types.

**Supplementary Table 25**. Results of gene expression analysis for “Survived” vs. “Died” comparison among “pathogen-negative” samples for all tested cell types.

**Supplementary Table 26**. Results of gene expression analysis for “CAP/HAP/NPC” (timepoint 1) vs. “no VAP” (timepoint 2) comparison for serial samples for all tested cell types.

**Supplementary Table 27**. Results of gene expression analysis for “CAP/HAP/NPC” (timepoint 1) vs. “VAP” (timepoint 2) comparison for serial samples for all tested cell types.

**Supplementary Table 28**. Results of gene expression analysis for “Future no VAP” (timepoint 1) vs. “Future VAP” (timepoint 1) comparison for serial samples for all tested cell types.

**Supplementary Table 29**. Results of gene expression analysis for “VAP will be cured within 7 days” vs. “VAP will not be cured within 7 days” comparison for all tested cell types.

**Supplementary Table 30**. Results of gene expression analysis for samples from patients “< 65 years old” vs. “≥ 65 years old” for all tested cell types.

**Supplementary Table 31**. Results of gene expression analysis for samples from male patients vs. female for all tested cell types.

**Supplementary Table 32**. Results of gene expression analysis for samples from immunocompetent patients vs. immunocompromised for all tested cell types.

**Supplementary Table 33**. Results of gene expression analysis for samples performed when patients received “≤ 600 cumulative steroid dose” (median dose) vs. “> 600” for all tested cell types.

**Supplementary Table 34**. Results of gene expression analysis for samples performed when patients spent “< 2 days on mechanical ventilation” (first quartile) vs. “> 16” (fourth quartile) for all tested cell types.

**Supplementary Table 35**. List of genes encoding secreted proteins, obtained from Human Protein Atlas on 11th of June, 2025.

**Supplementary Table 36**. Performance of our Geneformer-based models in predicting sample-level pathogen classification, and cure, measured by area under receiver operating characteristic curve (AUROC) and area under precision-recall curve (AUPRC).

**Supplementary Table 37**. Pathogen groupings and per-pathogen CFU cutoffs for all pathogens detected in the cohort.

**Supplementary Table 38**. Numbers of samples for each value of each categorical clinical covariate tested in MOFA association analysis.

